# Multiplexed, multimodal profiling of the intracellular activity, interactions, and druggability of protein variants using LABEL-seq

**DOI:** 10.1101/2024.04.19.590094

**Authors:** Jessica J. Simon, Douglas M. Fowler, Dustin J. Maly

## Abstract

Multiplexed assays of variant effect are powerful tools for assessing the impact of protein sequence variation, but are limited to measuring a single protein property and often rely on indirect readouts of intracellular protein function. Here, we developed LAbeling with Barcodes and Enrichment for biochemicaL analysis by sequencing (LABEL-seq), a platform for the multimodal profiling of thousands of protein variants in cultured human cells. Multimodal measurement of ∼20,000 variant effects for ∼1,600 BRaf variants using LABEL-seq revealed that variation at positions that are frequently mutated in cancer had minimal effects on folding and intracellular abundance but could dramatically alter activity, protein-protein interactions, and druggability. Integrative analysis of our multimodal measurements identified networks of positions with similar roles in regulating BRaf’s signaling properties and enabled predictive modeling of variant effects on complex processes such as cell proliferation and small molecule-promoted degradation. LABEL-seq provides a scalable approach for the direct measurement of multiple biochemical effects of protein variants in their native cellular context, yielding insight into protein function, disease mechanisms, and druggability.

## Introduction

A protein’s sequence defines its properties, including biophysical characteristics, intracellular function, pathogenicity, druggability and drug resistance. Measurement of protein variant effects has provided valuable insight into mechanisms of protein allostery^1, 2^ and regulation^3^, drug resistance and sensitivity^4, 5^, molecular evolution^6, 7^, and the effects of germline and somatic mutations^8–10^. Probing how protein sequence variation affects protein properties has also guided design and engineering efforts^11^ as well as enabled the development and benchmarking of computational models of protein stability and structure. Multiplexed assays of variant effect (MAVEs) are powerful tools for characterizing the consequences of protein sequence variation because they leverage high-throughput DNA sequencing to enable the measurement of thousands of protein variants in a single experiment. However, while first-generation MAVEs have enabled measurement of the properties and functions of millions of variants of 100s of proteins^12^, they are generally confined to measuring a single protein property or function, are crafted for a specific protein or related family members, are difficult to scale, and may divorce the protein from its cellular context. Many MAVEs also measure aspects of protein function indirectly; for example, using cell proliferation as a proxy for enzymatic activity. Thus, a general, scalable platform for measuring the effects of protein sequence variation using direct biochemical readouts of protein properties in a human intracellular environment is urgently needed.

The multi-domain kinase BRaf is emblematic of the need for such a platform. BRaf plays a central role in oncogenic signaling and >250 somatic mutations have been identified so far in The Cancer Genome Atlas (TCGA), ClinVar, and COSMIC databases^13–15^. While numerous elegant biochemical and cellular studies have been performed, they have focused on a limited subset of variants and do not provide a comprehensive view of the effects of BRaf sequence variation^16, 17^. Thus, our understanding of the mechanistic basis of BRaf regulation and signaling, the role that BRaf wild type (WT) and specific variants play in disease-relevant signaling, and the druggability of clinically-observed variants remains limited. In unstimulated cells, BRaf WT exists mainly within an autoinhibited complex. Upon activation, BRaf undergoes a series of large conformational changes and reprogramed protein-protein interactions that tune its ability to modulate downstream signaling. Consequently, BRaf sequence variation can affect multiple coupled intra- and intermolecular interactions. Therefore, to fully understand BRaf’s function, multiple biochemical properties must be measured in the native human cell signaling environment where all regulatory components and interaction partners are present. Applying existing MAVEs to measure all of these properties would require considerable effort and the creation of a new assay for each biochemical property.

Thus, we developed LAbeling with Barcodes and Enrichment for biochemicaL analysis by sequencing (LABEL-seq), which enables multiplexed, multimodal biochemical profiling of thousands of protein variants in cultured human cells. LABEL-seq can be used to comprehensively profile intracellular abundance, activity, protein-protein interactions and other properties of a protein variant library. Applying LABEL-seq to ∼1,600 BRaf variants revealed that substitutions at positions where somatic mutations are frequently observed minimally affected folding and intracellular abundance. However, mutations at many of these same positions had dramatic effects on activity, interactions with multiple signaling partners, and druggability. Integrative analysis of our multimodal variant effect data revealed groups of positions that play similar mechanistic roles in modulating BRaf’s regulation and function and illustrated how LABEL-seq-derived biochemical measurements can enable predictive modeling of the gain-of-function properties that BRaf variants acquire to promote growth factor-independent proliferation. Finally, LABEL-seq revealed that, surprisingly, 22.9% of BRaf variants were sensitized and 13.5% were resistant to small molecule-promoted degradation. These biochemical variant effects were integrated to construct a model that predicted sensitivity and resistance to small-molecule protein degradation, shedding light on the factors influencing druggability. In total, applying LABEL-seq to ∼1,600 BRaf variants generated ∼20,000 variant effect measurements of abundance, activity, protein-protein interactions, and druggability. Thus, LABEL-seq is a platform technology for measuring protein variant effects applicable to multiple biochemical properties and functions, facilitating a deeper understanding of the complex cellular behavior of proteins.

## Results

### LABEL-seq, a modular platform for multiplexed profiling of protein variants

To facilitate direct biochemical analyses of protein variant properties in a human cellular environment, we developed LABEL-seq, a modular platform that utilizes the intracellular self-assembly of proteins with nucleic acid barcodes to form complexes that remain stably associated following cell lysis and biochemical enrichment (Fig. 1a). LABEL-seq enables multiplexed profiling of the biochemical properties of protein variants in an intracellular environment by quantifying barcodes using high-throughput DNA sequencing following biochemical enrichment. Multiple biochemical properties of the same protein variant can be measured by tailoring which protein component self-assembles with a nucleic acid barcode and by modifying the enrichment strategy.

**Fig. 1.**
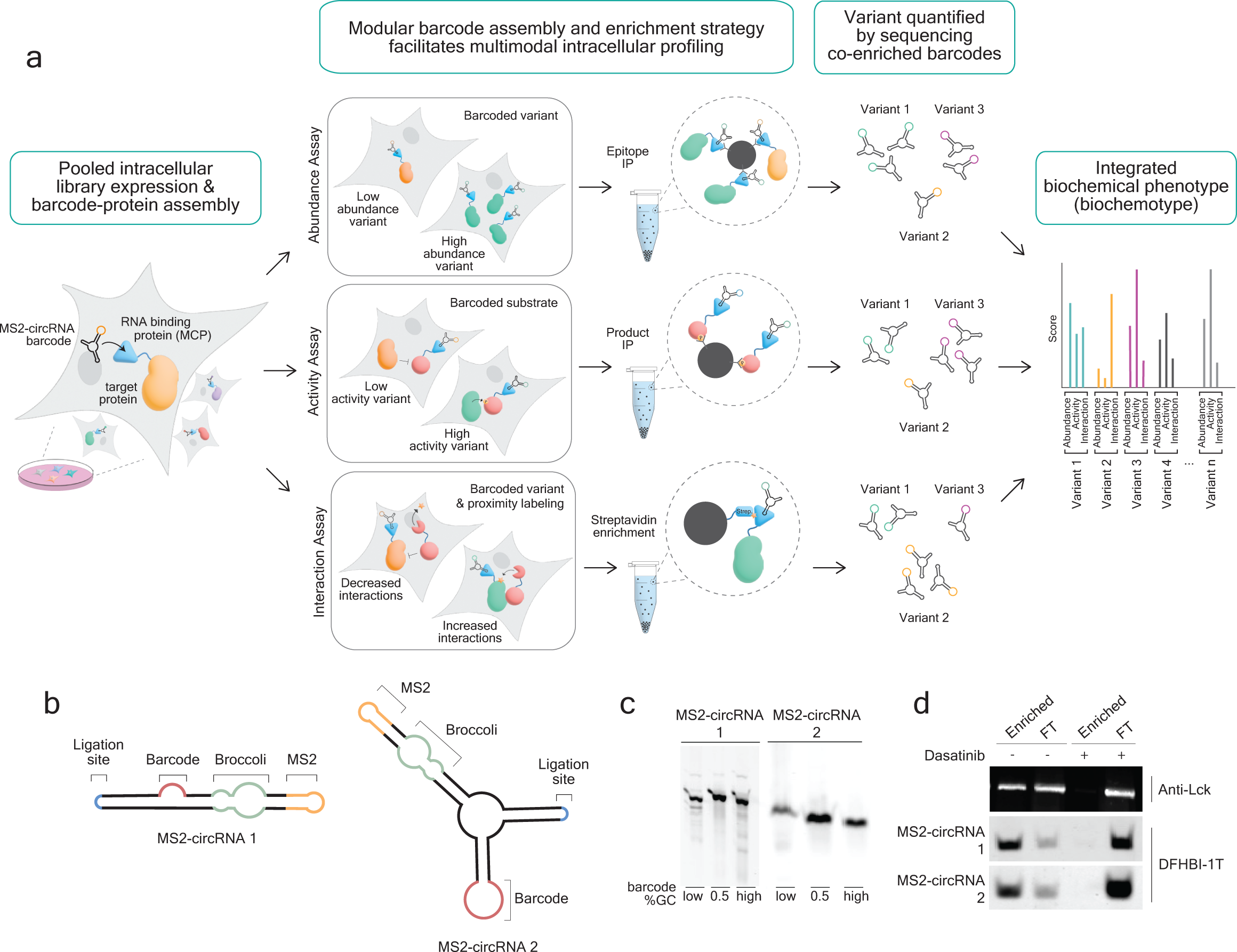
Overview of LABEL-seq and development of the RNA and protein components. **a,** Schematic depicting the LABEL-seq workflow. Immunoprecipitation (IP). **b,** Two MS2-circRNA architectures containing MS2 RNA hairpins and degenerate 16 nucleotide barcodes. **c,** In-gel fluorescence of total RNA extracted from cells expressing MS2-circRNAs 1 or 2 and stained with DFHBI-1T. **d**, Enrichment of tdMCP-Lck protein fusions complexed with MS2-circRNAs 1 or 2 with the immobilized ATP-competitive inhibitor dasatinib. FT = flow through of enrichment. + = presence of free dasatinib competitor.

As the barcoding component of LABEL-seq, we developed nucleic acids that are (1) transcriptionally generated, (2) stable to cellular and biochemical enrichment conditions, (3) tolerant of the addition of a 15-20 nucleotide barcode and, most importantly, (4) contain the MS2 RNA stem loop, which stably self-assembles with the viral MS2 coat protein (MCP)^18^. Because RNA can be transcriptionally generated, we selected single-stranded RNA as the genetically-encoded barcoding component. Due to the general instability of RNA, we pursued a ribozyme-assisted circRNA architecture that has been shown to be highly stable in an intracellular environment^19, 20^. We used RNAfold to guide the placement of a 19 nucleotide MS2 hairpin and a 16 nucleotide barcode within two different circRNA architectures (“MS2-circRNAs”, Fig. 1b and Extended Data Fig. 1a,b) and identified seven MS2-circRNA designs where barcode insertion and replacement of a stem loop with the MS2 hairpin were predicted to minimally alter the folding of the terminal ribozymes required for circularization. We found that most of these designs expressed at comparably high levels to the parent circRNAs and were minimally affected by barcode identity (Fig. 1c and Extended Data Fig. 1c). MS2-circRNAs 1 and 2, which represent the two different circRNA architectures, were selected for further investigation.

We fused four different variants of MCP (Extended Data Fig. 1d) to the N-terminus of the tyrosine kinase Lck and tested each for their ability to enrich co-expressed MS2-circRNA. We found that all four MCP-Lck fusions efficiently enriched co-expressed MS2-circRNA 2 when captured with the immobilized kinase inhibitor dasatinib, but that undetectable levels of each MCP-Lck fusion and MS2-circRNA 2 were captured in the presence of a free dasatinib competitor (Extended Data Fig. 1e). We selected a tandem MCP dimer (tdMCP) that contains a deletion that prevents higher-order oligomerization^21–23^ and an affinity-enhancing mutation (V29I)^21, 23^ for further characterization and observed that this tdMCP variant efficiently co-enriched MS2-circRNAs 1 and 2 (Fig. 1d).

### Self-assembled tdMCP:MS2-circRNA complexes maintain fidelity between protein variant-barcode associations

We engineered a HEK293 landing pad line^24^, circRNA-Protein Landing Pad (RPLP), that facilitates efficient integration of a single protein variant and a single MS2-circRNA barcode sequence per cell in a pooled format (Extended Data Fig. 2a). The RPLP line contains a Bxb1 recombination site at a single genomic locus that is flanked by Tet-inducible and U6 promoters (Extended Data Fig. 2b), which allows a transgenic cassette containing a protein variant and MS2-circRNA barcode sequence pair to be expressed only after recombination. We next recombined the RPLP with plasmids containing a MS2-circRNA 1b library (∼20,000 unique barcode sequences) and a Flag-tagged tdMCP-BRaf fusion. Immunoprecipitation of tdMCP-BRaf from cell lysate revealed barcode sequence counts were robustly correlated between two replicate enrichments (Pearson’s R^2^ = 0.93, Fig. 2a). Pulldowns from as few as 25 cells per barcode provided high correlation between barcode sequence quantifications (Pearson’s R^2^ = 0.80; Extended Data Fig. 2c) as did immunoprecipitations of a different protein kinase target, tdMCP-BTK, co-expressed with a library of MS2-circRNA barcode sequences (Pearson’s R^2^ = 0.98, Extended Data Fig 2d).

**Fig. 2.**
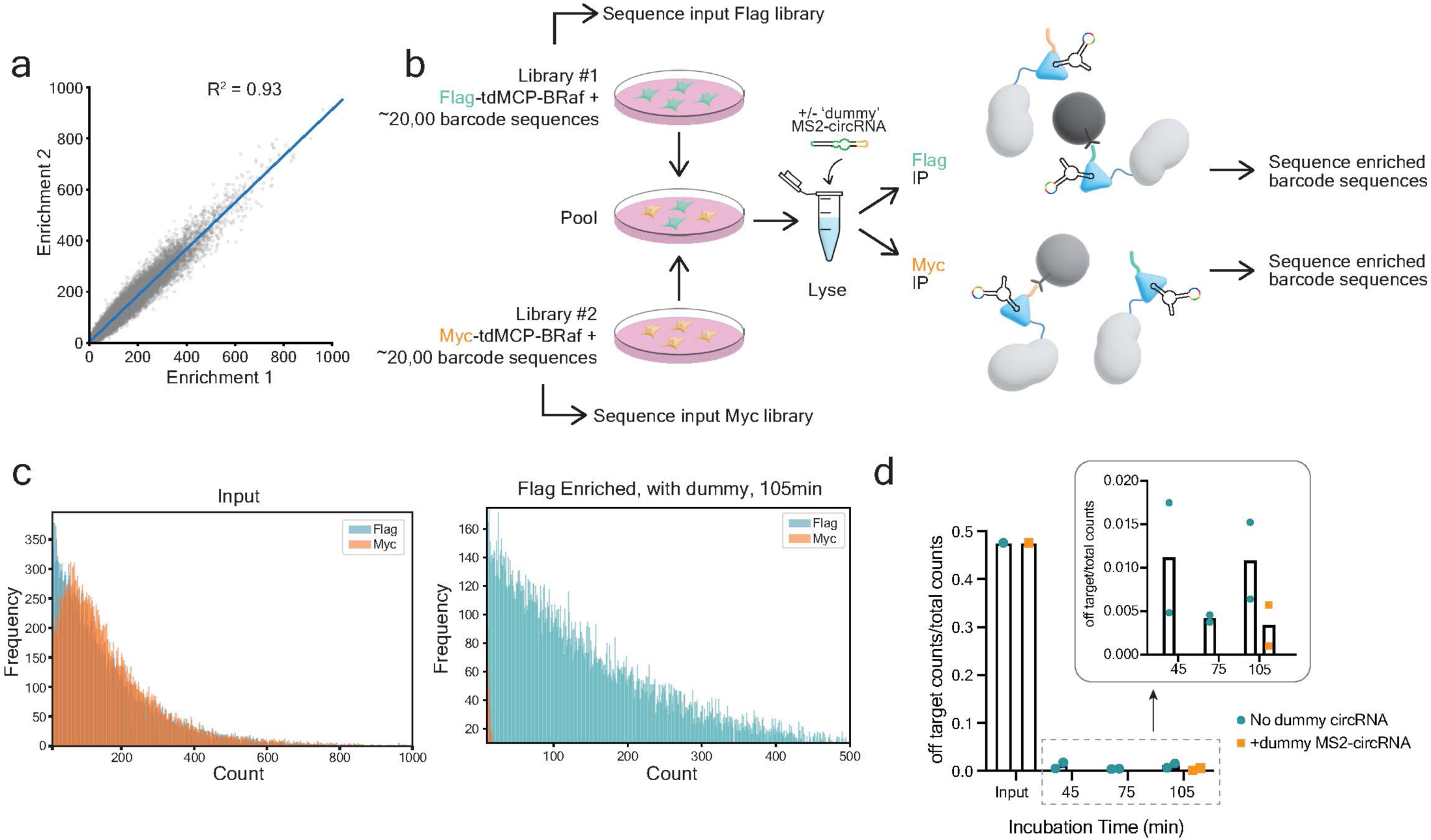
Protein:MS2-circRNA complexes are stable during biochemical enrichment. **a,** Sequenced barcode counts derived from two independent Flag immunoprecipitations (IP) of Flag-tdMCP-BRaf co-expressed with a library of MS2-circRNA 1b barcode sequences (Pearson’s R^2^ = 0.93). **b,** Experimental workflow of the barcode sequence misassociation experiment. Dummy MS2-circRNA does not contain a barcode or the priming site required for reverse transcription. **c,** Results from the experiment shown in **b**. Relative frequencies of barcode sequence counts matched to Flag-tdMCP-BRaf and Myc-tdMCP-BRaf libraries in unenriched lysate (left) and after 105 min Flag immunoprecipitation performed with lysate that was generated with dummy MS2-circRNA in the lysis buffer (right). **d,** Quantification of the data shown in **c**. The y-axis is the fraction of barcode sequence counts encoding the Myc-tagged library (n=2 expression, lysis, enrichment, and sequencing replicates).

Each barcode sequence must remain stably associated with its cognate protein following lysis and during enrichment because dissociation and reassociation of protein:MS2-circRNA complexes would diminish dynamic range. Therefore, we measured the extent of MS2-circRNA barcode sequence misassociation following cell lysis and a prolonged enrichment (Fig. 2b). Into the RPLP, we separately integrated a MS2-circRNA 1b (Extended Data Fig. 1b) library containing ∼20,000 barcode sequences encoding a Flag-tagged tdMCP-BRaf fusion and an MS2-circRNA 1b library containing ∼20,000 orthogonal barcode sequences encoding a Myc-tagged tdMCP-BRaf fusion. After recombination, we combined the two orthogonally barcoded libraries, performed parallel Myc and Flag immunoprecipitations, and sequenced the barcodes from each enrichment. Only ∼1% of barcode sequences enriched in the Flag immunoprecipitation were associated with the Myc library, despite equal input frequencies (Fig. 2c,d). Notably, the fraction of barcode sequences associated with the Myc library did not vary at different immunoprecipitation timepoints, suggesting that minimal barcode exchange occurred during enrichment. Furthermore, we observed that when an excess of an MS2-circRNA that does not contain a barcode and cannot be reverse transcribed (“dummy” MS2-circRNA, Extended Data Fig. 2e) was added to the lysis buffer, only ∼0.3% of the quantified barcode sequences encoded Myc after a 105 min enrichment (Fig. 2c,d). Comparably low levels of off-target Flag-tdMCP-BRaf barcode sequences were enriched following immunoprecipitation of Myc-tdMCP-BRaf (Fig. 2d and Extended Data Fig. 2f,g). Together, libraries of MS2-circRNAs efficiently express and assemble with co-expressed tdMCP-protein fusions and protein variant-barcode sequence associations are maintained during prolonged enrichments, thereby enabling the analysis of variant libraries at scale.

### Profiling of protein variant abundance with LABEL-seq

We selected the multi-domain kinase BRaf to demonstrate the multimodal profiling capabilities of LABEL-seq due to the complexity of its regulation, BRaf’s ability to interact with multiple signaling partners, and the prevalence of disease-associated mutations^25^. We first confirmed that an N-terminal tdMCP fusion^26, 27^ of BRaf behaves like untagged BRaf. We observed that previously characterized BRaf variants promoted expected levels of pMek (Fig. 3a), co-expression of CRaf enhanced tdMCP-BRaf’s activation of downstream signaling (Extended Data Fig. 3a), and that tdMCP-BRaf possessed similar levels of phosphorylation at a regulatory site as untagged BRaf. (Extended Data Fig. 3b). In order to understand how sites that have been observed to be mutated in cancer affect the signaling properties of BRaf, we generated a library of single amino acid variants at all positions in BRaf that contain two or more missense mutations in TCGA and COSMIC databases (Supplementary Table 1). Two positions (R509 and I666) that contain biochemically well-characterized variants were also varied, for a total of 80 positions that are distributed throughout the entire protein (Fig. 3b and Extended Data Fig. 3c). Long read sequencing of the resulting ∼1,600 member tdMCP-BRaf variant library indicated each variant was represented by ∼30 unique MS2-circRNA 1b barcode sequences (Extended Data Fig. 3d), which was recombined into the RPLP.

**Fig. 3.**
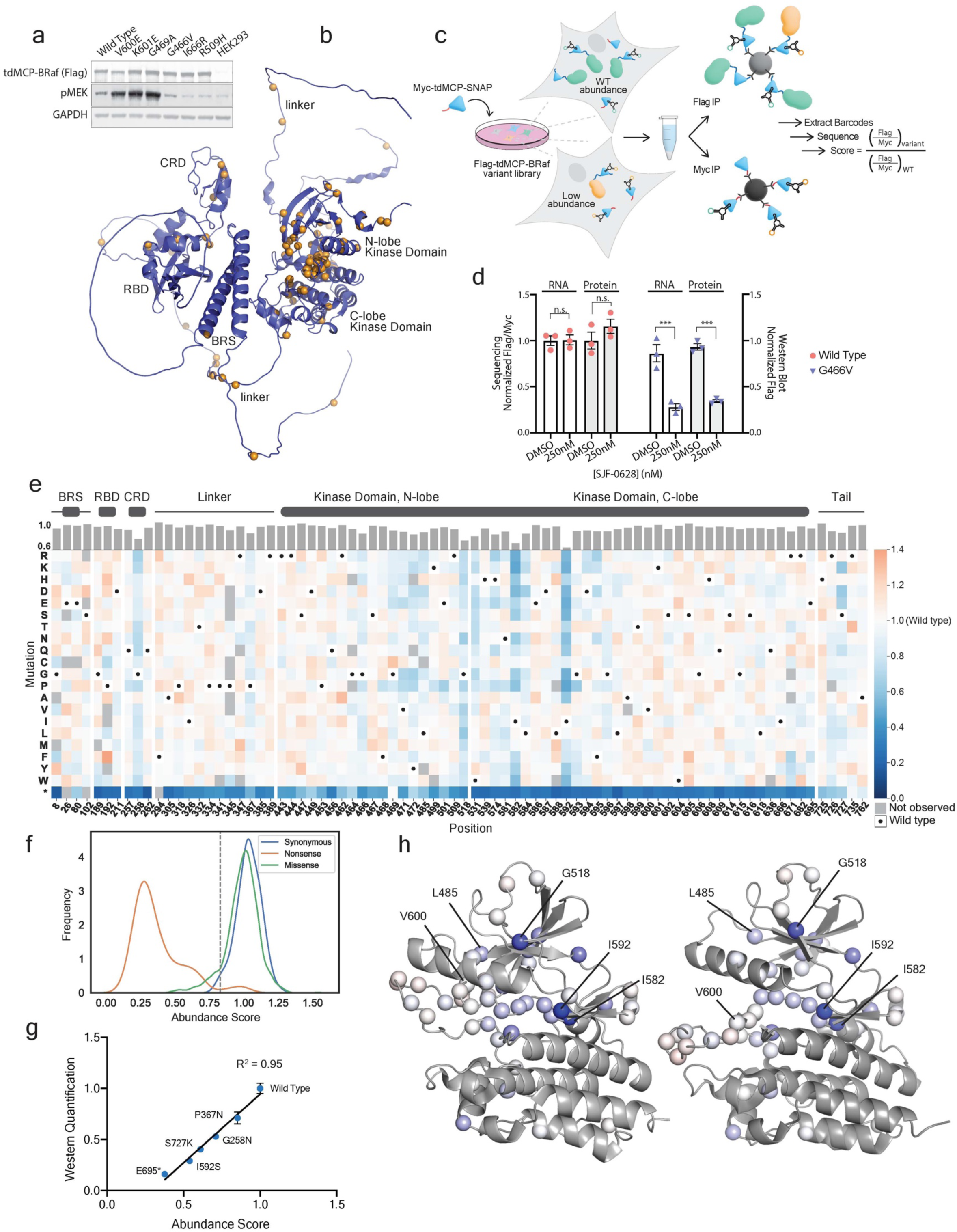
Multiplexed abundance measurements reveal BRaf’s tolerance for substitution at positions where somatic mutations are frequently observed. **a,** Western blot analysis of HEK293 RPLP cells stably expressing tdMCP-BRaf variants. **b,** AlphaFold2 model (AF-15056-F1) of full-length human BRaf. Varied positions in the untagged and tdMCP-BRaf variant libraries are shown as orange spheres. **c,** Schematic depicting the LABEL-seq abundance assay in which a tdMCP standard (Myc-tdMCP-SNAP) is co-expressed with a tdMCP-variant library. Abundance scores are WT-normalized ratios of variant barcode sequence counts derived from parallel Flag and Myc immunoprecipitations (IP). **d,** Comparison of abundance measurements obtained by western blotting (right, y-axis = ratio of Flag signal to Myc signal normalized to Flag-tdMCP-BRaf WT-expressing cells treated with DMSO) or by quantifying co-enriched MS2-circRNA barcode sequences with high-throughput sequencing. (left, y-axis = the ratio of barcode sequence counts derived from parallel Flag and Myc enrichments normalized to Flag-tdMCP-BRaf WT-expressing cells treated with DMSO). HEK293T cells co-expressing MS2-circRNAs, the Myc-tdMCP-SNAP standard, and Flag-tdMCP-BRaf WT (pink circles) or G446V (blue triangles) were treated with DMSO or the variant-selective degrader SJF0628 for 24 h prior to lysis. The average of 4 barcode sequence ratios per condition is shown per replicate (marked by symbol). Error bars = standard error of the mean (SEM) of replicate scores. Westerns are shown in Extended Data 3f. **e,** Sequence-abundance map for BRaf variants in the tdMCP-BRaf library. The abundance scores shown are the average of two replicates, each involving an independent Myc-tdMCP-SNAP standard transduction, cell lysis, parallel Flag and Myc immunoprecipitations, and quantification by high-throughput sequencing. Black dots in the map indicate the WT amino acid and gray tiles indicate missing data. Position-averaged abundance scores are shown in the bar graph above the heatmap. **f,** LABEL-seq abundance score distributions for all synonymous WT and nonsynonymous BRaf variants. The gray dashed line indicates the abundance score value (>2 standard deviations of the synonymous distribution lower than WT) we defined as decreased abundance. **g,** Individually assessed variant abundances measured by western blotting (n=3) compared to abundance scores determined with the LABEL-seq abundance assay for a panel of BRaf variants (Pearson’s R^2^ = 0.95). Error bars = SEM. **h,** Position-averaged abundance scores for each position in the tdMCP-BRaf variant library (represented as spheres) projected onto the structures of (left) autoinhibited (PDB ID: 6NYB) and (right) active BRaf (PDB ID: 4MNE). The shade of blue indicates position-averaged abundance score: white = 1, darkest blue = 0.72.

Quantitative measurements of protein variant abundance are important because all proteins must express at high enough levels to perform their cellular function^10, 28, 29^ and many complex biochemical properties, such as activity and protein-protein interactions, cannot be fully understood in the absence of protein stability information^30^. To enable the measurement of a range of protein abundances, we developed a LABEL-seq abundance assay that involves co-expressing Flag-tdMCP-tagged protein variants with a Myc-tdMCP standard, which serves as an intracellular competitor for MS2-circRNAs (Fig. 3c), and calculating the ratio of barcode sequence counts obtained from parallel Flag and Myc immunoprecipitations. This ratiometric quantification provides a direct readout of abundance that is not dependent on the relative intracellular expression levels of MS2-circRNAs and protein variants. Prior to abundance profiling of our tdMCP-BRaf variant library, we confirmed that the sequencing-derived abundance scores of cells co-expressing the Myc-tdMCP standard with tdMCP-BRaf WT or G466V, and treated with a variant-selective BRaf PrOteolysis TArgeting Chimera (PROTAC)^31^, reflected levels determined by western blot analysis (Fig. 3d and Extended Data Fig. 3e).

We next used the LABEL-seq abundance assay to profile the ∼1600 BRaf variants in our library (Fig. 3e and Supplementary Table 2). We found that abundance scores were highly correlated (Pearson’s R^2^=0.75, Extended Data Fig. 3f) between biological replicates, and that the synonymous WT and nonsense score distributions were well separated (Fig. 3f). As expected, we observed that proline and charged amino acid substitutions most negatively impact BRaf abundance (Extended Data Fig. 3g)^29^. Notably, the LABEL-seq-calculated abundance scores of individual BRaf variants representing a range of expression levels were highly correlated with the levels of variants that were not fused to tdMCP assessed by western blot quantification (Pearson’s R^2^=0.95, Fig. 3g). Thus, LABEL-seq can accurately and precisely measure protein variant abundance at scale.

Our abundance profiling shows that the 80 profiled positions of BRaf were highly tolerant of substitution, with 86.8% of missense variants demonstrating WT-like abundance (Fig. 3f and Extended Data Fig. 3g). The permissiveness of these 80 positions to undergo substitution without affecting abundance is notable based upon previous observations where up to 50% of substitutions are not tolerated in some proteins^28^, but is consistent with their location on BRaf. Profiled regions of BRaf include the linker connecting the cysteine-rich domain (CRD) to the kinase domain, the kinase domain’s phosphate-binding loop (P-loop), activation loop, and C-terminal tail, which are all flexible and structurally dynamic (Extended Data Fig. 3h,i). In comparing the structures of autoinhibited and activated BRaf, many of the positions profiled also undergo large conformational transitions between regulatory states^32, 33^(Fig. 3h). Calculation of position-averaged abundance scores showed that select positions are relatively intolerant to substitution (Fig. 3h). These positions are situated in regions that are comparably less conformationally dynamic than other profiled positions, with four of the five lowest average abundance scores being constituents of β-strands despite only 12.5% of all profiled positions being classified as β-stranded (Extended Data Fig. 3i). The two positions that were least tolerant to substitution, I582 and I592, are located on adjacent β-strands, make hydrophobic contacts, and are structurally invariant in the autoinhibited and active forms of BRaf (Fig. 3h and Extended Data Fig. 3j). Our results are consistent with systematic analyses showing that disease-relevant mutations commonly alter the biochemical properties of a protein without dramatically affecting its folding and stability,^34,35^ and suggest that most sites of somatic cancer mutations in BRaf can readily accumulate gain-of-function activities because substitutions at these positions have a minimal effect on abundance.

### Profiling of intracellular BRaf variant activity with LABEL-seq

We next developed a LABEL-seq assay that provides quantitative measurements of intracellular enzymatic activity by co-expressing untagged protein variants and their encoding MS2-circRNAs with tdMCP-fused substrates and selectively enriching products (Fig. 4a). Because Raf activates Mek and the activation loop of Erk is only phosphorylated by Mek (Fig. 4b), we used Flag-tdMCP-Erk2 as the substrate (pErk reporter) and anti-pErk enrichment as a readout of Raf variant activity. We first confirmed that the pErk reporter was efficiently enriched from lysate (Extended Data Fig. 4a) and that enriched barcode sequence frequencies were higher from cells expressing hyperactive BRaf V600E compared to BRaf WT (Fig. 4c). Next, we used the LABEL-seq activity assay to profile an untagged BRaf variant library (Extended Data Fig. 3d) in RPLP cells co-overexpressing CRaf because WT and many BRaf variants signal as heterodimers (Fig. 4d,e)^34^. Activity scores were calculated using the ratio of WT-normalized barcode sequence counts derived from parallel pErk and Flag immunoprecipitations (Fig. 4d and Supplementary Table 2). Activity scores from two biological replicates were well correlated (Pearson’s R^2^ = 0.80, Fig. 4f and Extended Data Fig. 4b,c) and BRaf variants previously shown to promote elevated pErk^34–36^ had significantly higher activity scores than WT (Extended Data Fig. 4c). Activity scores of individual BRaf variants that represent a range of activity levels were also highly correlated with pErk levels quantified by western blotting (Pearson’s R^2^ = 0.95, Fig. 4g).

**Fig. 4.**
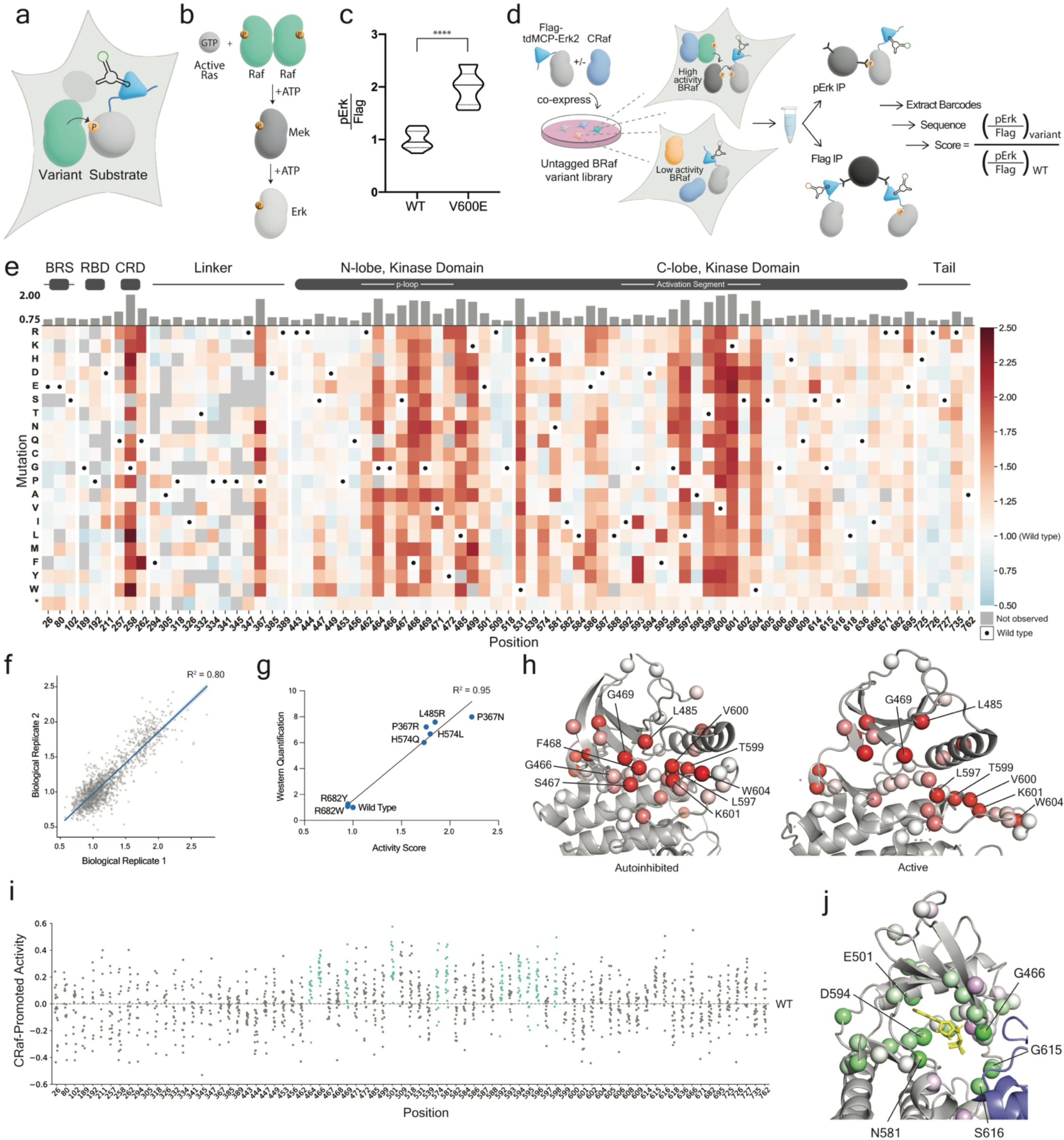
A pErk reporter enables profiling of intracellular BRaf variant activity. **a,** General schematic of the LABEL-seq activity assay. **b,** The canonical Raf-Mek-Erk signaling pathway. **c,** Activities (represented as the ratio of pErk and Flag barcode sequence counts) of BRaf WT and V600E in HEK293T cells co-expressing Flag-tdMCP-Erk2 (pErk reporter) and MS2-circRNAs. Cells were lysed, parallel Flag and pErk immunoprecipitations were performed, and the ratio of pErk/Flag counts for each co-enriched MS2-circRNA barcode sequence was calculated. Center line = mean; dotted lines = inner quartiles. n=10-11 barcodes per variant. ****P<0.001. **d,** Schematic of the LABEL-seq assay for the multiplexed measurement of untagged BRaf variant activity with the pErk reporter. **e,** Sequence-activity map for variants in the untagged BRaf library co-expressed with CRaf. The activity scores shown are the average of two replicates, each involving independent cell culturing, lysis, parallel pErk and Flag immunoprecipitations, and quantification with high-throughput sequencing. Black dots in the map indicate the WT amino acid and gray tiles indicate missing data. Position-averaged activity scores are shown in the bar graph above the heatmap. The average activity score at each position is shown in the bar graph above the heatmap. **f,** Scatterplot showing activity score correlations between the two independent replicates shown in **e** (Pearson’s R^2^ = 0.80). **g,** Individually assessed pErk levels measured by western blotting (n=3) compared to activity scores determined with the LABEL-seq activity assay for a panel of BRaf variants (Pearson’s R^2^ = 0.95). Error = SEM. **h,** The number of activating variants at each position in the untagged BRaf variant library (represented as spheres) projected onto the (left) inactive (PDB ID: 6NYB) and (right) active conformations (PDB ID: 4MNE) of BRaf’s kinase domain. The number of activating variants at a position is represented by the shade of red: white = 0 activating variants, brightest red = 19 activating variants. G466, S467, and F468 are not resolved in the active conformation of BRaf’s kinase domain. **i,** CRaf-promoted, BRaf variant activity at each position in the untagged BRaf variant library. CRaf-promoted activity scores were calculated by determining the difference between BRaf variant activity scores, which were each normalized to BRaf WT in the absence or presence of co-expressed CRaf. Positions highly conserved amongst protein kinases are shown in green. Gray dashed line indicates the WT score (0). Positive and negative values indicate more or less activity promoted by CRaf, respectively. **j,** Position-averaged CRaf-promoted activity scores projected onto the structure of the autoinhibited BRaf:Mek complex (PDB ID: 6NYB) bound to ATP-γ-S. Purple spheres are scores less than 0 (decreased activity with CRaf) and green spheres are scores greater than 0 (increased activity with CRaf).

We classified variants as activating or inactivating (Extended Data Fig. 4d,e), and found that 32.8% and 2.3% were activating and inactivating, respectively (Extended Data Fig. 4e,f). The small percentage of inactivating variants and the observation that positions involved in the formation of active Raf dimers possessed the lowest average activity scores (Extended Data Fig. 4g) is consistent with BRaf WT existing primarily as an autoinhibited monomer in unstimulated HEK293 cells. Observed activating variants are broadly distributed across BRaf, with a number of positions containing multiple activating mutations.

Because BRaf WT mainly exists in an inactive monomeric state stabilized by a network of interactions in unstimulated cells, we reasoned that positions participating in autoinhibitory interactions should possess multiple activating mutations. We identified 17 potentially autoinhibitory positions where >75% of substitutions were classified as activating (Supplementary Table 3). Two of these positions–G258 and P367–are in BRaf’s N-terminal CRD and RBD domains, respectively, and substitutions at these positions likely disrupt autoinhibitory protein-protein interactions (Extended Data Fig. 4h,i). Nine potentially autoinhibitory positions form a spatial cluster in the kinase domain of autoinhibited but not active BRaf (Fig. 4h). In the autoinhibited conformation, the side chains of four of these positions in the activation loop (L597, T599, V600 and K601) form interactions with three positions in the P-loop (S467, F468, and G469) and L485. The side chain of W604 does not form obvious contacts with these positions but appears to be part of this autoinhibitory network based on its proximity and high percentage of activating mutations. Our data are consistent with previous speculation based on structural studies that eight of these nine positions participate in stabilizing an “autoinhibitory turn”^36–38^, and suggest that this region of BRaf’s kinase domain contains a finely tuned autoinhibitory interaction network that can readily be disrupted through mutation to achieve increased BRaf activity.

To examine the effects of heterodimerization on BRaf variant activity, we repeated the LABEL-seq activity assay in the absence of overexpressed CRaf, which provided broadly similar patterns of activating and inactivating mutations (Extended Data Fig. 5a-d), and calculated the difference in activity score with and without CRaf (“CRaf-promoted activity”, Figure. 4i). We observed that seven of the ten positions with the highest average CRaf-promoted activity scores are highly conserved across the human kinome^39^ and are largely clustered in and around the ATP-binding pocket (Fig. 4j and Extended Data Fig. 5e). Among these highly conserved positions, mutations at G466, N581, D594, and G596 have been previously demonstrated to impair BRaf’s kinase activity^34, 36^ while retaining the ability to increase cellular pErk levels through heterodimerization and transactivation of CRaf^34^. Two other positions with the highest average CRaf-promoted activity, G615 and S616, are not conserved but are positioned on the BRaf:Mek interface (Fig. 4j), suggesting that substitutions at these positions disrupt BRaf’s interaction with its substrate but not its ability to activate CRaf in trans. Together, our data show that disruption of catalytic machinery within the ATP-binding site or prevention of Mek binding is a general mechanism for increasing BRaf’s reliance on CRaf for promoting downstream signaling.

### Proximity labeling-based profiling of intracellular protein-protein interactions with LABEL-seq

Raf activity is modulated by a number of intracellular protein-protein interactions. In the absence of active Ras, BRaf WT is stabilized as an autoinhibited monomer in a complex with its substrate, Mek, and two 14-3-3 proteins. Upon Ras activation, BRaf undergoes large conformational changes and forms active homodimers or heterodimers with CRaf. To gain a comprehensive understanding of how sequence variation affects BRaf’s intracellular interactions, we developed a LABEL-seq interaction assay using TurboID proximity labeling and streptavidin enrichment (Fig. 5a and Extended Data 6a-e). We first validated that TurobID could reliably report on BRaf interactions by co-expressing a TurboID-Mek1 fusion with tdMCP-BRaf WT or I666R, a BRaf:Mek-disrupting variant, and treating cells with biotin. We observed that tdMCP-BRaf I666R showed expected reduced biotinylation levels relative to WT (Fig. 5b) and that streptavidin enrichment followed by barcode sequence quantification recapitulated these differences (Fig. 5c). Both tdMCP-BRaf variants showed markedly higher signals than an unrelated cytosolic kinase (tdMCP-BTK), which was used as an estimate of non-specific labeling (Fig. 5c). We further found that the LABEL-seq interaction assay provided expected increases in tdMCP-BRaf WT labeling by TurboID-CRaf or TurboID-BRaf in cells treated with a dimer-promoting inhibitor (Fig. 5d-f and Extended Data Fig. 6f).

**Fig. 5.**
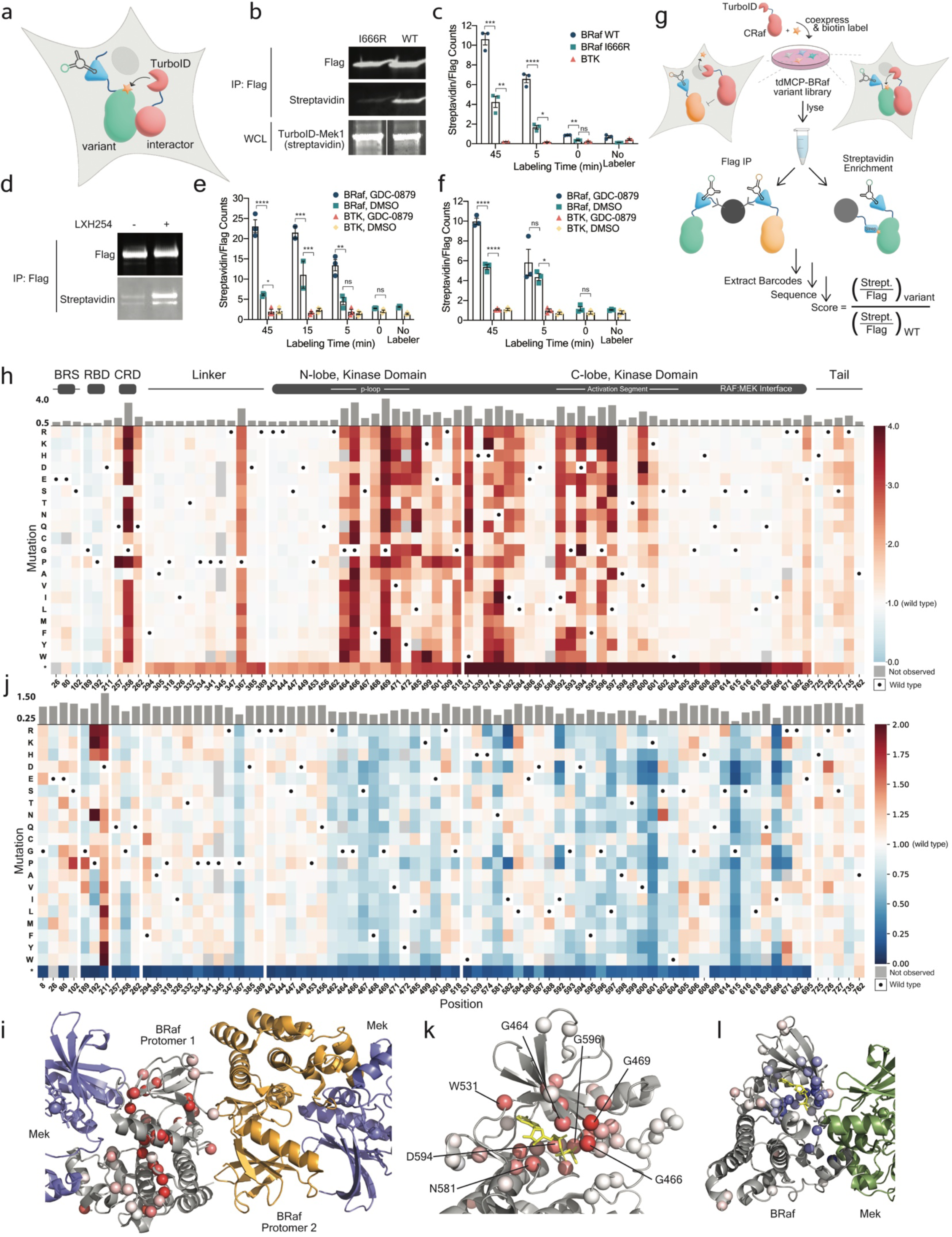
LABEL-seq measurements of the intracellular interactions of BRaf variants with CRaf and Mek1. **a,** General schematic of the proximity labeling-based LABEL-seq interaction assay. **b,** Biotinylation levels of Flag-tdMCP-BRaf I666R (left) and WT (right) in HEK293 cells co-expressing TurboID-Mek1 and labeled with biotin for 90 min. **c**, Biotinylation levels (represented as the ratio of streptavidin and Flag barcode sequence counts) of Flag-tdMCP-BRaf WT (blue circles), I666R (teal squares) and background control Flag-tdMCP-BTK (pink triangles) in HEK293T cells co-expressing TurboID-Mek1 and MS2-circRNAs, and treated with biotin for 0, 5, or 45 min. Following biotin treatment, cells were lysed, pooled, parallel streptavidin and Flag enrichments were performed, co-enriched barcodes were quantified by high-throughput sequencing, and the ratio of streptavidin and Flag counts for each co-enriched MS2-circRNA barcode sequence was calculated (n=3 barcodes/treatment, mean +/- SEM, *P<0.0332, **P<0.0021, ***P<0.0002, ****P<0.0001). **d**, Biotinylation levels of Flag-tdMCP-BRaf WT in HEK293T cells co-expressing TurboID-CRaf, treated with DMSO (left) or type 2 inhibitor LXH-254 (right) for 4 h, and then labeled with biotin for 90 min. **e,** Biotinylation levels (represented as the ratio of streptavidin and Flag barcode sequence counts) of Flag-tdMCP-BRaf WT and Flag-tdMCP-BTK in HEK293T cells co-expressing TurboID-CRaf and MS2-circRNAs, which were treated DMSO or GDC-0879 for 2 h followed by labeling with biotin for 0, 5, 15, or 45 min. Following biotin addition, cells were lysed, parallel streptavidin and Flag enrichments were performed, and the ratio of streptavidin and Flag counts for each co-enriched MS2-circRNA barcode sequence was calculated using high-throughput sequencing. **f,** Biotinylation levels (represented as the ratio of streptavidin and Flag barcode sequence counts) of Flag-tdMCP-BRaf WT and Flag-tdMCP-BTK in HEK293T cells co-expressing TurboID-BRaf and MS2-circRNAs, which were treated DMSO or GDC-0879 for 2 h followed by labeling biotin for 0, 5, or 45 min. Following biotin addition, cells were treated as described in **e**. For **e**,**f** (n=2-3 barcodes/treatment, mean +/- SEM, *P<0.0332, **P<0.0021, ***P<0.0002, ****P<0.0001) **g**, Schematic of the LABEL-seq assay for the multiplexed measurement of intracellular BRaf variant interactions. **h,** Sequence-CRaf interaction map for variants in the tdMCP-BRaf library co-expressed with TurboID-CRaf and labeled with biotin for 5 min. The interaction scores shown are the average of two replicates, each involving an independent cell culturing, biotin labeling, parallel streptavidin and Flag enrichments, and quantification with high-throughput sequencing. Black dots in the map indicate the WT amino acid and gray tiles indicate missing data. Position-averaged CRaf interaction scores are shown in the bar graph above the heatmap. **i,** The number of variants at each position that were classified as increased CRaf interaction in **h** projected onto a structure of BRaf’s kinase domain within the active BRaf dimer complex (PDB: 4MNE). White = 0 increased CRaf interaction variants, brightest red = 19 increased CRaf interaction variants. **j,** Sequence-Mek1 interaction map for variants in the tdMCP-BRaf library co-expressed with TurboID-Mek1 and labeled with biotin for 5 min. The Mek1 interaction scores shown are the average of two replicates, each involving an independent transfection, biotin labeling, parallel streptavidin and Flag enrichments, and quantification with high-throughput sequencing. Black dots indicate the WT amino acid and gray tiles indicate missing data. Position-averaged Mek1 interaction scores are shown in the bar graph above the heatmap. **k,** Position-averaged CRaf interaction scores for each position in the tdMCP-BRaf variant library (represented as spheres) projected onto a structure of autoinhibited BRaf bound to ATP-γ-S (PDB ID: 6NYB). The shade of red indicates the position-averaged CRaf interaction score: white = 1, brightest red > 3.2. **l,** Position-averaged Mek1 interaction scores projected onto the structure of the autoinhibited BRaf complex (PDB ID: 6NYB). The shade of blue indicates the position-averaged Mek1 interaction score: white = 1, darkest blue < 0.5.

We first used our LABEL-seq interaction assay to profile BRaf:CRaf heterodimerization by co-expressing TurboID-CRaf with the tdMCP-BRaf variant library, labeling cells with biotin, and calculating CRaf interaction scores (Fig. 5g,h and Supplementary Table 1). We observed that interaction scores were well correlated between biological replicates (Pearson’s R^2^ = 0.96, Extended Data 7b) and between 5 min and 45 min labeling times (Pearson’s R^2^ = 0.86, Extended Data Fig. 7a). We further found that the interaction score of BRaf WT was markedly higher than the non-specific labeling control, tdMCP-BTK, and that BRaf variants previously shown to exhibit elevated heterodimerization with CRaf^34^ had interaction scores significantly above BRaf WT (Extended Data Fig. 7c). We next classified variants as forming increased or decreased CRaf interactions (Extended Data Fig. 7d,e). Consistent with BRaf WT mainly existing as an autoinhibited monomer in unstimulated HEK293 cells, only 5.1% of BRaf variants were classified as decreased interaction (Extended Data 7e). There are a number of positions distributed throughout BRaf where a majority of variants were classified as increased CRaf interaction (Fig. 5i), suggesting that substitutions at these positions disrupt specific interactions that stabilize monomeric BRaf. Notably, nonsense mutations that truncate BRaf C-terminal to the RBD and CRD provided some of the highest interaction scores, confirming that BRaf’s CRD can act in trans with CRaf’s kinase domain^40^. Given that BRaf WT signals as a dimer, we were curious whether most mutations that promote CRaf heterodimerization are also activating. Although we did not find a strong correlation between activity and CRaf interaction scores (Extended Data Fig. 7f), 82% (406) of the 493 missense variants classified as activating were also classified as increased CRaf interaction. Together, our CRaf interaction data support the model that BRaf WT exists primarily as an autoinhibited monomer, which somatic mutations or upstream signaling events are capable of releasing to allow CRaf heterodimerization.

We next profiled the interactions of BRaf with its substrate by co-expressing TurboID-Mek1 with the tdMCP-BRaf variant library, labeling cells with biotin (Fig. 5j and Supplementary Table 1), and calculating Mek1 interaction scores (Pearson’s R^2^ = 0.80, Extended Data Fig. 7g-i). Interaction scores for BRaf I666R and the non-specific labeling control were both significantly lower than BRaf WT (Extended Data Fig. 7j), and Mek1 interaction scores were consistent with levels of Mek1 co-IP’d with individual BRaf variants (Extended Data Fig. 7k). Unlike CRaf, a larger percentage of mutations decreased (28.4%) rather than increased (6.0%) BRaf’s interaction with Mek1 (Extended Data Fig. 7h), consistent with autoinhibited BRaf WT existing in a stable complex with Mek. As expected, the lowest interaction scores were for nonsense variants that truncated BRaf prior to the disordered C-terminal tail of the kinase domain (Fig. 5j and Extended Data Fig. 7i) and the five sites directly lining the BRaf:Mek interface. Notably, 84% of missense variants not on the BRaf:Mek interface that were classified as decreased Mek interaction were also classified as increased CRaf interaction, suggesting that weakening the interaction with Mek allows heterodimerization with CRaf.

To obtain a structural understanding of which positions contribute to BRaf’s interactions with CRaf and Mek1, we projected position-averaged interaction scores onto the structure of autoinhibited BRaf bound to an ATP analogue (Fig. 5k,l). Consistent with ATP interactions stabilizing monomeric BRaf^37^, we observed that most positions that line the ATP-binding pocket possessed high position-averaged CRaf interaction scores. Intriguingly, positions with low average Mek1 interaction scores also line the ATP-binding pocket, suggesting that disruption of BRaf’s interactions with ATP weakens a stable BRaf:Mek autoinhibitory complex (Fig. 5l and Extended Data Fig. 7l,m). Together, our data highlight a unique feature of BRaf regulation where ATP, Mek, and monomeric BRaf form a stable autoinhibitory complex that can readily be disrupted through mutation to allow BRaf:CRaf heterodimerization.

### Hierarchical clustering identifies groups of BRaf positions with similar biochemotypes

Our profiling data indicate substitutions at certain positions in BRaf result in dramatic changes to multiple biochemical properties, but we were unable to identify positions with similar functions or gain a mechanistic understanding of the positional effects of substitution using pairwise analyses (Extended Data Fig. 8a). Therefore, we performed hierarchical clustering based on the six biochemical properties measured with LABEL-seq to group positions into nine distinct biochemical phenotypes (biochemotypes) (Fig. 6a,b). Clusters 7–9, which encompassed 64% of profiled positions, were characterized by a biochemotype in which most variants exhibited modest effects on the six measured biochemical properties. In contrast, cluster 1 positions are defined by a biochemotype where mutations resulted in some of the largest decreases in Mek interactions and largest increases in activity and CRaf heterodimerization, with minimal CRaf-promoted activity (Fig. 6a,c). One cluster 1 position–P367–is proximal to the autoinhibitory phosphorylation site, S365^41^, and sits within a 14-3-3 binding pocket in the autoinhibited BRaf complex (Extended Data Fig. 4i). Substitutions at P367 likely disrupt the autoinhibited BRaf complex and allow formation of active Raf dimers based on the known binding specificity of 14-3-3 proteins^42^ and the biochemotype of this position. The six other cluster 1 positions are located in two different regions of BRaf’s ATP-binding site (Fig. 6c). Four of these positions (F468, L485, L597 and V600) are components of the autoinhibitory turn (Fig. 4h) and are spatially clustered in the autoinhibited but not the active form of BRaf, suggesting that mutations at these positions promote increased activity and CRaf heterodimerization by disrupting autoinhibitory interactions. Despite their disparate locations, positions in cluster 1 appear to share a common functional role in stabilizing the autoinhibited state, which can be readily disrupted by mutations without compromising the ability of BRaf variants to form catalytically active Raf dimers.

**Fig. 6.**
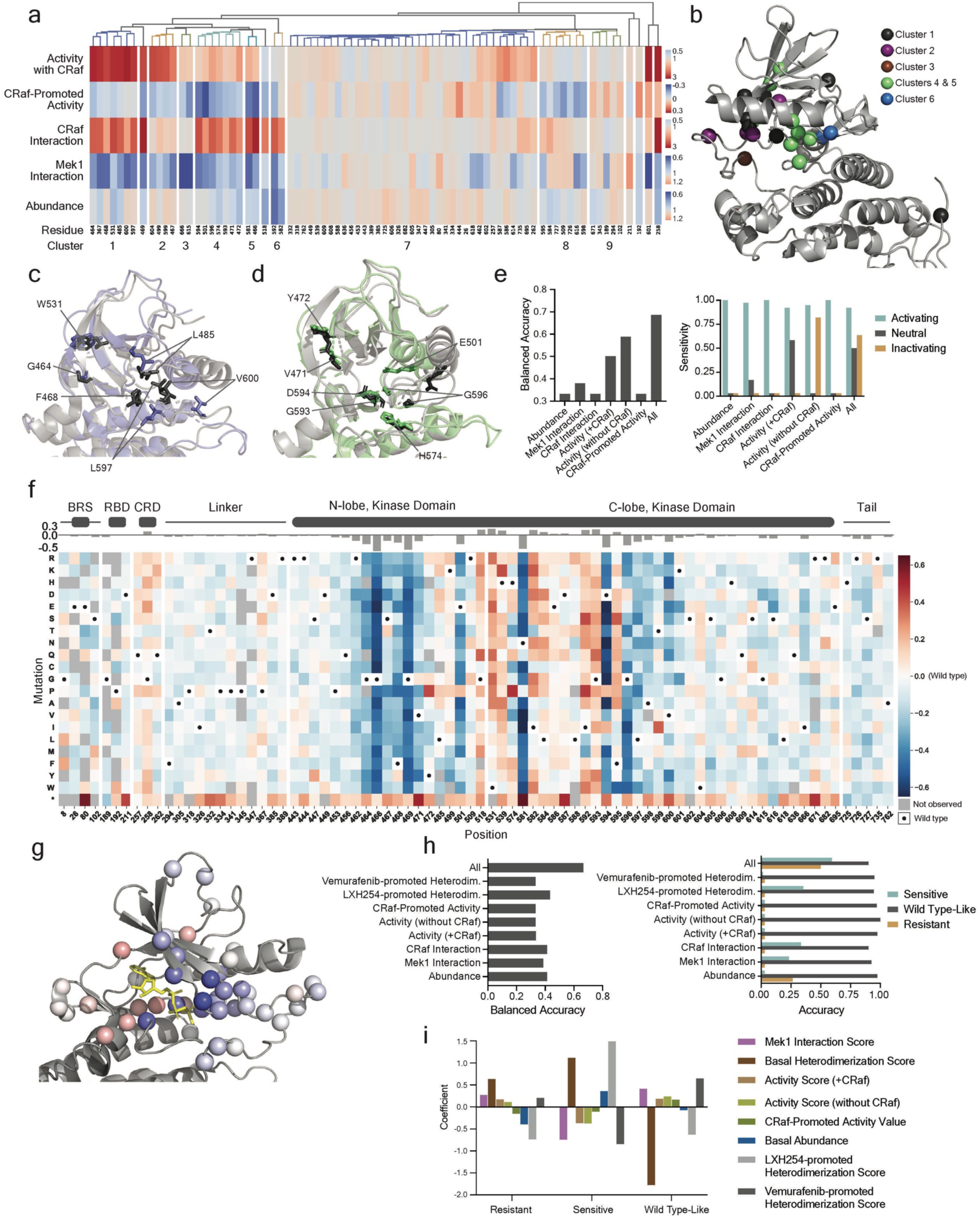
Integration of multimodal measurements. **a,** Heatmap, with the dendrogram shown above, showing hierarchical clustering of positions within BRaf based on six biochemical properties measured with LABEL-seq. Clustering was performed using the average distance metric, which grouped positions based on similarities in response to specific substitutions. Clusters of positions, which were selected based on the dendrogram, are numbered. The heatmap color indicates the average score for each biochemical property at each position. The color scale for each biochemical property is shown on the far right. **b,** Positions within clusters 1-6 mapped onto the kinase domain of the autoinhibited BRaf complex (PDB ID: 6NYB). **c,** Alignment of BRaf’s ATP-binding pocket in the active (purple, PDB ID: 4MNE) and autoinhibited (gray, PDB ID: 6NYB) conformations, with cluster 1 positions shown as sticks. F468 is not resolved in the active conformation of BRaf. **d,** Alignment of BRaf’s ATP-binding pocket in the active (green, PDB ID: 4MNE) and autoinhibited (gray, PDB ID: 6NYB) conformations, with cluster 4 positions shown as sticks. **e,** Performance of our logistic model predicting variant-driven proliferation of Ba/F3 and MCF10A cells in the absence of growth factor. (left) Balanced accuracy for models that were fit by using a single property versus all properties. (right) Model accuracy for each classification. **f,** Sequence-degradation map for BRaf variants in the tdMCP-BRaf library treated with 2.5 μM SJF0628. Degradation score = [abundance score for a BRaf variant at 2.5 μM SJF0628 - abundance score for a variant in DMSO]/abundance score for a variant in DMSO. The degradation scores shown are the average of two replicates, each involving an independent cellular treatment with SJF0628, parallel Flag and Myc immunoprecipitations, and quantification by high-throughput sequencing. Black dots in the map indicate the WT amino acid and gray tiles indicate missing data. The position-averaged degradation score is shown in the bar graph above the heatmap. **g,** Position-averaged degradation scores for each position in the tdMCP-BRaf variant library (represented as spheres) projected onto the structure of autoinhibited BRaf bound to ATP-γ-S (PDB ID: 6NYB). The color indicates the position-averaged degradation score: white = 0, darkest blue < -0.5, brightest red > 0.18. **h,** Performance of a logistic model predicting SJF0628 sensitivity. (left) Balanced accuracy for models fit using a single property versus all properties. (right) Model accuracy for each classification. **i,** Coefficients for each property in the most accurate model.

Cluster 2 positions, proximal to cluster 1 autoinhibitory turn positions (Extended Data Fig. 8b), exhibited the biochemotype of increased activity and decreased Mek interactions upon substitution like cluster 1, but without the accompanying strong increase in CRaf heterodimerization. The ability of mutations in cluster 2 to increase kinase activity and decrease Mek interactions suggest that these positions also play a role in stabilizing autoinhibited BRaf. However, mutations at cluster 2 positions appear to compromise BRaf’s strong association with CRaf once autoinhibition is released. Positions in clusters 4 and 5 demonstrated a biochemotype where substitutions strongly promoted CRaf heterodimerization and decreased Mek interactions, but, in contrast to clusters 1 and 2, only modestly increased Raf activity, which was highly CRaf dependent. Cluster 4 and 5 positions are located deep within BRaf’s active site (Fig. 6b,d) and form a contiguous spatial network in the active form of BRaf (Fig. 6d). Additionally, nearly all positions in clusters 4 and 5 are highly conserved amongst protein kinases^39, 43^ and many make contacts with ATP^37^ or are part of the kinase catalytic machinery (Extended Data Fig. 5e). The biochemotype of clusters 4 and 5 suggest that these positions participate in the stabilization of autoinhibited BRaf. However, mutations at these positions likely impair BRaf’s kinase activity, rendering cluster 4 and 5 variants dependent on their ability to form strong CRaf heterodimers for Mek activation. Together, our multimodal analysis reveals that a number of positions in BRaf that are frequently mutated in cancer participate in finely tuned autoinhibitory interactions that most mutations are capable of destabilizing. The signaling properties of these BRaf variants with disrupted autoinhibition can be classified into several distinct biochemotypes based on their location.

### LABEL-seq-guided prediction of a variant-promoted cellular phenotype

We next determined whether LABEL-seq data could be used to predict which biochemical properties of BRaf variants are capable of driving complex phenotypes such as cell proliferation. Here, we trained logistic models using previously collected data that classified 65 BRaf variants as activating, inactivating, or neutral based on their ability to promote the growth factor-independent proliferation of Ba/F3 and MCF10A cells^44^. Models that used only one biochemical property had modest classification accuracy (balanced accuracy: 0.33-0.57l; Fig. 6e). However, a model that used all six biochemical properties measured with LABEL-seq provided the highest accuracy (balanced accuracy = 0.68) and, critically, was the only model capable of correctly identifying BRaf variants of all three classifications (activating, incactivating, or neutral; Fig. 6e). We applied this six-property classifier to the 239 somatic mutations cataloged in TCGA, COSMIC, and ClinVar and for which we obtained LABEL-seq data, and found that 40.6% were predicted to be activating and 14.2% were predicted to be inactivating (Supplemental Table 3). Notably, 92% of somatic variants previously determined to be activating based on increased intracellular pErk levels or growth factor-independent cell proliferation in separate analyses were correctly classified by our model as activating (Supplemental Table 1). Thus, measuring multiple biochemical properties for the same variant, enabled by LABEL-seq, facilitated prediction of a complex cellular phenotype.

### Measurement and modeling of small molecule-promoted BRaf degradation

We next explored whether LABEL-seq data could be used to provide insight into another poorly understood cellular process: small molecule-promoted protein degradation^45^. Specifically, we investigated the Vemurafenib-based PROTAC SJF0628 (Extended Data Fig. 9a), which promotes selective degradation of several BRaf variants^31^. We first used the LABEL-seq abundance assay to profile the degradation of the tdMCP-BRaf variant library across a range of SJF0628 concentrations (Fig. 6f and Supplementary Table 1) and calculated degradation scores for all ∼1,600 variants (Extended Data Fig. 9b,c). As expected, BRaf WT was largely resistant to degradation by SJF0628 (Supplementary Table 2) but a number of variants, including previously characterized SJF0628-sensitive variants^31^ (Extended Data Fig. 9d), showed dose-dependent decreases in abundance. Additionally, LABEL-seq-derived degradation scores were consistent with levels of protein degradation quantified by western blot (Extended Data Fig. 9c). We classified variants as sensitized or resistant to degradation at the highest concentration of SJF0628 tested and observed that 22.9% of variants were sensitized and 13.5% were resistant (Extended Data Fig. 9e and Supplementary Table 1). Among the 239 somatic mutations for which we measured degradation scores and that are listed in TCGA, COSMIC, or ClinVar, we found that 32.3% were sensitized to degradation relative to BRaf WT and 9.2% were resistant. Notably, variants at the same position were generally either all sensitized or all resistant, suggesting that susceptibility to degradation by SJF0628 is largely defined by the position of a mutation rather than its identity.

We explored the structural basis for the location dependence of BRaf variant PROTAC sensitivity by projecting position-averaged degradation scores onto the structure of Vemurafenib-bound BRaf (Extended Data Fig. 9f). Several positions within the Vemurafenib-binding pocket generally conferred resistance to degradation when substituted but, surprisingly, some of the most sensitive positions were located on nearby positions without an obvious structural rationale for either property. Although projecting position-averaged degradation scores onto the autoinhibited structure of BRaf bound to an ATP analogue revealed that substitutions at positions that make direct contacts with ATP were largely sensitizing (Fig. 6g), positions with the lowest average degradation scores did not fit within any of the biochemotypes we identified. Furthermore, none of the biochemical properties we measured correlated strongly (R^2^ = 0.051-0.36) with variant sensitivity (Extended Data Fig. 9g).

We performed logistic modeling to identify which biochemical properties of BRaf variants were most predictive of PROTAC sensitivity. To provide additional properties that relate to ATP-competitive inhibitor binding of BRaf, we used the LABEL-seq interaction assay to measure CRaf heterodimerization levels promoted by the type 1.5 inhibitor Vemurafenib and the type 2 inhibitor LXH254^46^ (Extended Data Fig. 10a,b). The resulting eight biochemical properties, along with a subset of our measured degradation classifications were used to train a logistic model to predict whether a BRaf variant was sensitized, resistant, or WT-like in its degradation response at the highest concentration of SJF0628. Models using any one property yielded modest classification accuracy (balanced accuracy = 0.33-0.43), but the model using all eight properties was most accurate (balanced accuracy = 0.67, Fig. 6h). Only the model incorporating all eight properties could accurately distinguish BRaf variants across all three classifications (Fig. 6h). The properties most heavily weighted in predicting variant sensitivity to SJF0628 were the CRaf interaction and Type 2 inhibitor (LXH254)-promoted heterodimerization scores (Fig. 6i), suggesting that dimerization is an important component of degradation. Low basal abundance and Type 2 inhibitor (LXH254)-promoted heterodimerization scores were the most predictive of whether a variant was resistant to SJF0628. Notably, our data also indicate that it is not a particular “active” conformation of a variant that is most predictive of SJF0628 degradation, as previously speculated^31^, but each variant’s overall capacity to be promoted into Raf dimers. The ability of LABEL-seq to provide multimodal profiling of biochemical properties facilitates the analysis of complex cellular processes that are otherwise difficult to understand using single measurements.

## Discussion

At-scale measurements of protein variant effects have provided valuable insights into numerous biological processes, facilitated advances in protein engineering and design, and deepened our understanding of drug sensitivity and resistance^4, 8, 12^. To expand the scope of protein targets and functional properties that can be profiled at scale, we developed LABEL-seq, a general, scalable platform that enables the multimodal analysis of protein variants in human cells using high-throughput DNA sequencing for quantification.

A key strength of LABEL-seq is its modularity, which enables the profiling of multiple properties for a common set of protein variants. Reconfiguration of the RNA and protein components of LABEL-seq and tailoring of the biochemical enrichment step to measure specific protein properties enabled profiling of the abundance, activity, protein-protein interactions, and drug sensitivity of a library of BRaf variants. The high precision and sensitivity of quantification, and the flexibility to incorporate diverse biochemical enrichment strategies into the LABEL-seq workflow suggest that it can be used to profile numerous additional protein properties and functions. Even intracellular assays that have not been previously employed for studying protein variant properties, such as our use of proximity labeling to quantify both high and low affinity intracellular protein-protein interactions, are compatible with LABEL-seq. Finally, since LABEL-seq only necessitates the fusion of an RNA-binding domain to a target protein of interest or its substrate, it has the potential to be applied to a wide range of proteins beyond the three (BRaf, BTK, and Lck) described in this study.

We used LABEL-seq for the multimodal profiling of BRaf positions frequently mutated in cancer, complementing and expanding upon previous cellular, biochemical, and structural characterization of individual BRaf variants^34, 47^. Notably, we found that most mutations at cancer-associated positions in BRaf do not compromise BRaf’s folding and intracellular abundance. We confirmed previous observations that mutations at many of these same positions can promote increased Raf activity and dimerization^16, 36, 37^. Our LABEL-seq profiling data suggest that many positions in BRaf are precisely tuned to maintain an autoinhibited state, which is disrupted by most sequence variations. Consequently, BRaf is finely poised to acquire mutations that lead to enhanced downstream oncogenic signaling, likely due, at least in part, to its ability to form functionally active dimers upon disruption of autoinhibition. As our profiling experiments were conducted in the absence of elevated levels of active Ras, the biochemical properties of some BRaf variants that are relevant to oncogenic signaling may have been missed. Because LABEL-seq assays can be performed under variable cellular conditions, such as growth factor simulation or the co-expression of an active Ras mutant, future profiling experiments that capture these properties are feasible. Furthermore, while our profiling studies were conducted in HEK293 cells, it should be straightforward to achieve single protein variant and single MS2-circRNA barcode sequence integration into other cell lines, which will allow LABEL-seq profiling in other cellular contexts.

Our ability to use the LABEL-seq abundance assay to measure SJF0628’s degradation of BRaf variants allowed us to conduct the first comprehensive analysis of how sequence variation affects a neosubstrate’s degradation by a PROTAC. Remarkably, we observed that substitutions at multiple positions in BRaf provided strong sensitization to degradation, with SJF0628 selectively degrading ∼30% of somatic variants cataloged in TCGA, ClinVar, and COSMIC relative to BRaf WT. By modeling the multiple biochemical measurements obtained with LABEL-seq, we were able to obtain insight into which properties sensitize a BRaf variant to degradation by SJF0628. Although no single biochemical property was dominant, the ability of a BRaf variant to be recruited into heterodimers with CRaf was the most predictive of sensitivity to SJF0628-mediated degradation. We also identified several positions in BRaf that contained multiple mutations that conferred resistance to PROTAC-mediated degradation, providing insights into resistance mechanisms that complement those revealed by a previous CRISPR-suppressor scan of the neosubstrates GSPT1 and RBM39^48^. Conducting additional LABEL-seq assays with PROTACs that incorporate different E3 ligase recruiters and/or alternative ATP-competitive inhibitors will enable determination of whether the patterns of sensitization and resistance we observed are common to all BRaf-targeting PROTACs or are specific to SJF0628, which contains a ligand for recruiting the E3 ligase von Hippel-Lindau linked to the type 1.5 inhibitor vemurafenib. Furthermore, incorporating additional biochemical properties into models that more accurately predict BRaf variant degradation susceptibility to different PROTACs should yield additional mechanistic insight into this process.

Overall, we envision LABEL-seq will find widespread use for the multimodal profiling of diverse proteins under different cellular conditions. The multi-dimensional profiling data obtained with this platform will be a valuable resource for understanding intracellular protein function and druggability, and, when integrated with MAVEs that assess composite behaviors such as transcriptional outputs and proliferation, should provide invaluable insight into protein properties that drive complex cellular processes.

## Acknowledgements

We thank David Shechner (UW, Seattle) for guidance on RNA barcode design and Madeline Walsh (UW, Seattle) for critical feedback on the manuscript. This work was supported by the NIH, grant no. R01GM145011 (NIGMS) and R01GM086858 (NIGMS) to D.J.M.

## Author contribution statement

J.J.S., D.M.F., and D.J.M. conceived of the work and wrote the manuscript. J.J.S. developed the LABEL-seq method, performed all profiling experiments and data analysis.

---

**Extended Data Fig. 1.**
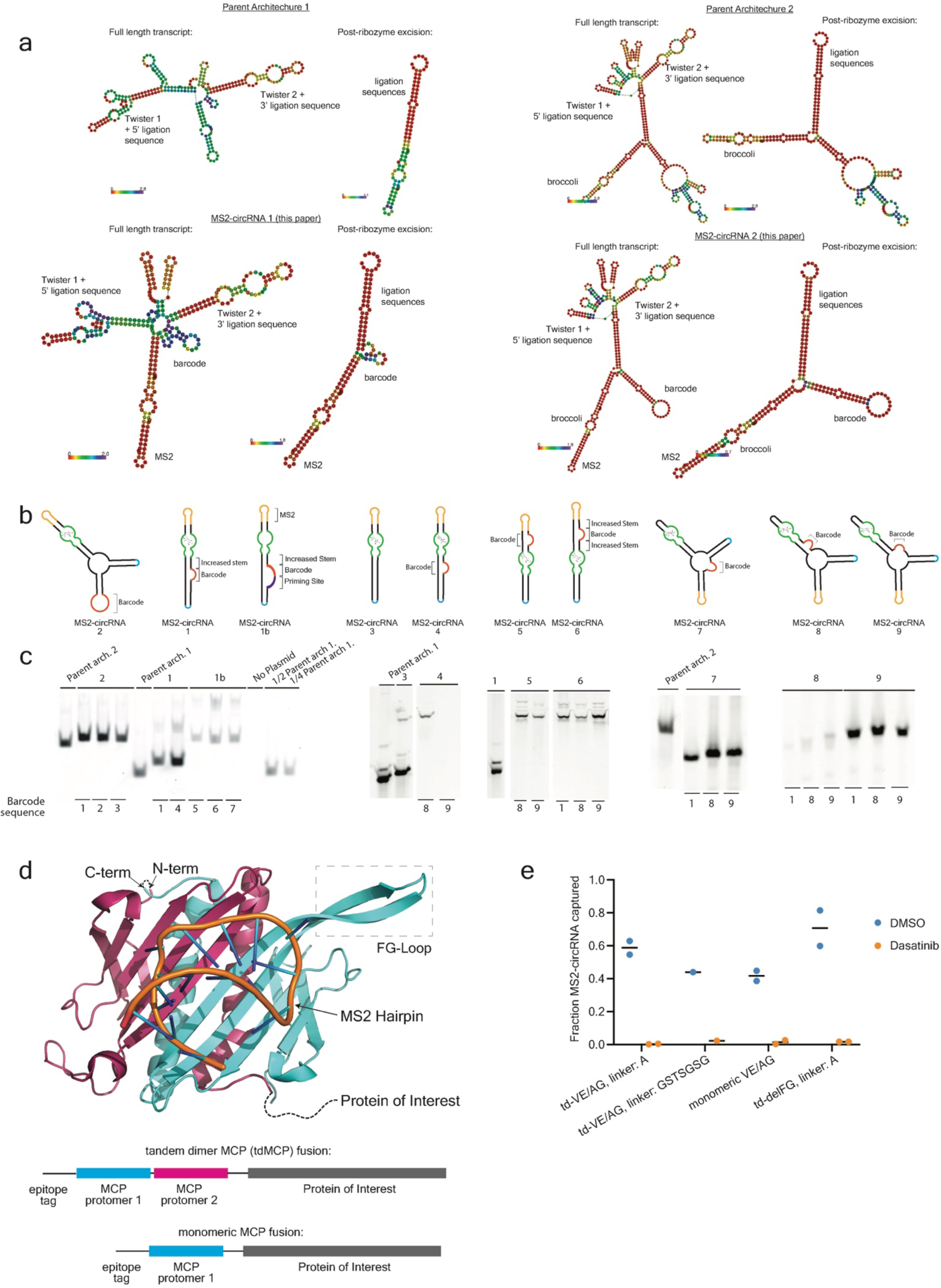
Design, expression, and characterization of MS2-circRNAs and MCP protein fusions. **a,** In-silico folding predictions for circRNA architectures pre- and post-ribozyme excision. Predictions were run using RNAfold and the colors indicate positional entropy values. Parent architectures do not contain MS2 and are the designs published by Litke and Jaffrey^19^ which correspond to addgene plasmids #124360 (left) and #124362 (right). **b,** MS2-circRNA designs that contain a 16 nucleotide barcode and 19 nucleotide MS2 hairpin. MS2-circRNA 1b contains an added priming site specifically for reverse transcription. **c,** In-gel DFHBI-1T fluorescence of total RNA extracted from cells expressing the MS2-circRNAs shown in b. 2-3 barcode sequences of varying GC content for each MS2-circRNA were tested. Gels 1, 4, and 5 were non-denaturing 6% acrylamide gels and gels 2 and 3 were denaturing 10% acrylamide gels. **d,** Structure (PDB ID: 2BU1) of the MCP dimer bound to an MS2 hairpin (top). Linear representations of monomeric and tandem-dimer (td) MCP protein fusions (bottom). In a tandem dimer, the N-terminus of one MCP protomer is linked to the C-terminus of a second MCP protomer by a short linker. **e,** Variants of MCP-Lck protein fusions were co-expressed with MS2-circRNA 2. Lck was enriched using dasatinib-conjugated agarose beads and the co-enriched MS2-circRNA was visualized by in-gel DFHBI-1T fluorescence. DMSO or a free dasatinib competitor was included during enrichment to probe for specific protein enrichment. td-indicates tandem dimer designs. VE/AG variants contain V75E and A81G mutations, which prevent MCP oligomerization. delFG variants possess a deletion of the FG loop (V67-A81), which also prevents MCP oligomerization. All MCP variants contain the high affinity V29I mutation. The linkers are positioned between MCP protomers. The delFG variant was used in all subsequent experiments and is referred to as ‘tdMCP’ throughout the manuscript.

**Extended Data Fig. 2.**
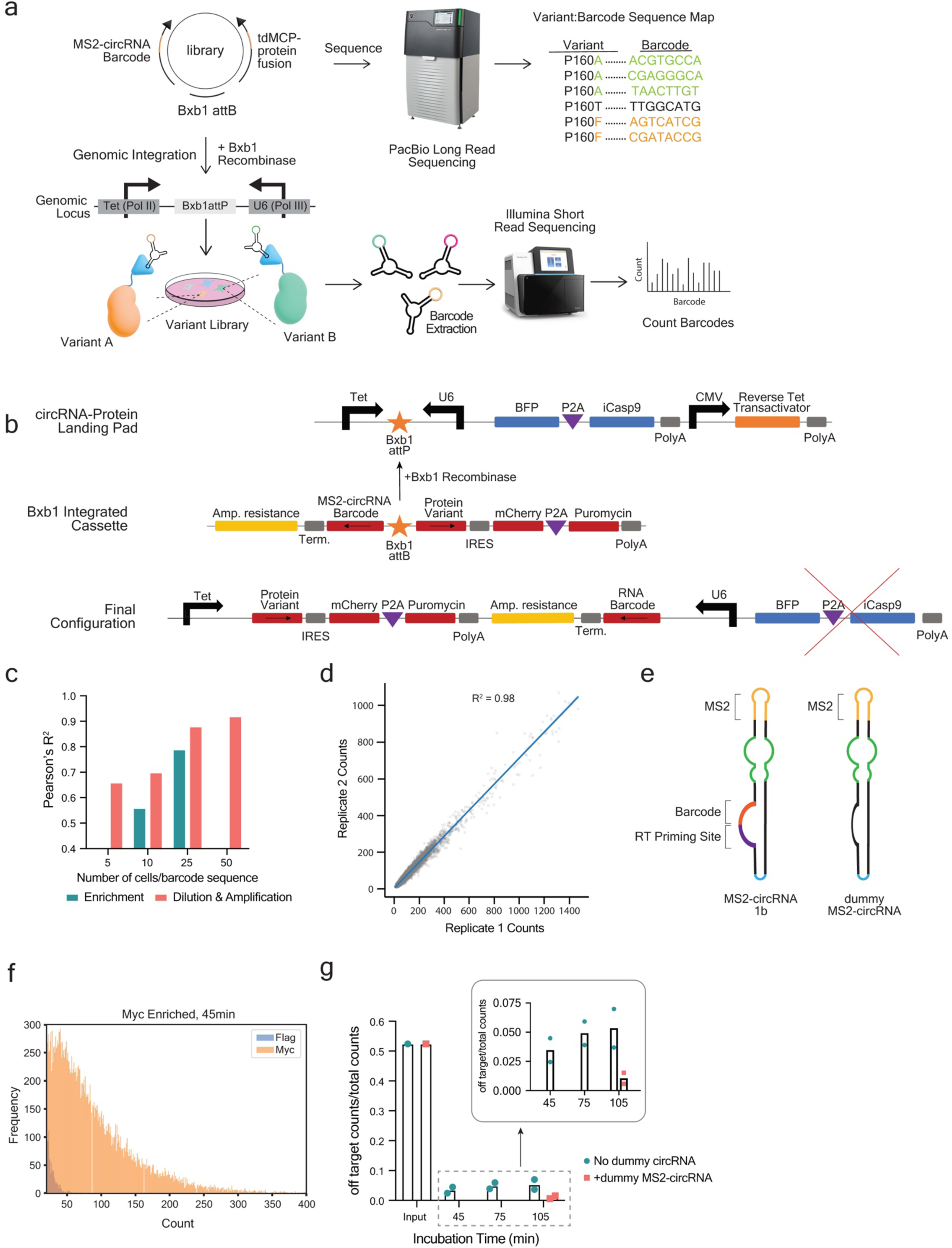
Engineering of a multiplexed MS2-circRNA barcoding platform. **a,** Subassembly, used to link protein variants to nucleotide barcodes, is achieved by sequencing the cloned plasmid library using long-read high-throughput sequencing. In parallel, the plasmid library is integrated into the RPLP to generate a cell line stably expressing tdMCP:MS2-circRNA complexes for use in biochemical experiments. **b,** Schematic of the RPLP cell line. Recombination of Bxb1 attB-containing plasmids into the RPLP stably integrates protein variants and MS2-circRNA barcodes under genomic transcriptional control. Selection for successful recombinants is run using AP1903 and/or puromycin. **c,** Parallel Flag immunoprecipitations were performed using serial dilutions of lysate generated from RPLP cells stably expressing Flag-tdMCP-BRaf WT and a library of 20,000 MS2-circRNA 1b barcode sequences. Pearson’s R^2^ correlation coefficients of barcode sequence counts are reported for replicate enrichments or RT-PCR amplifications. **d,** Sequenced barcode counts derived from two independent expressions and Flag immunoprecipitations of Flag-tdMCP-BTK co-expressed with a library of MS2-circRNA 2 barcode sequences. Pearson’s R^2^ = 0.98. **e,** Schematics of MS2-circRNA 1b, the architecture used to encode variants in the tdMCP- and untagged BRaf variant libraries, and dummy MS2-circRNA, which is included in the lysis buffer to minimize barcode sequence missassociation. **f,** Relative frequencies of barcode sequence counts matched to Myc-tdMCP-BRaf and Flag-tdMCP-BRaf libraries after 45 min Myc immunoprecipitation. **g,** Quantification of the data shown in **f**. The y-axis is the fraction of barcode sequence counts encoding the Flag-tagged library (n=2 expression, lysis, enrichment, and sequencing replicates).

**Extended Data Fig. 3.**
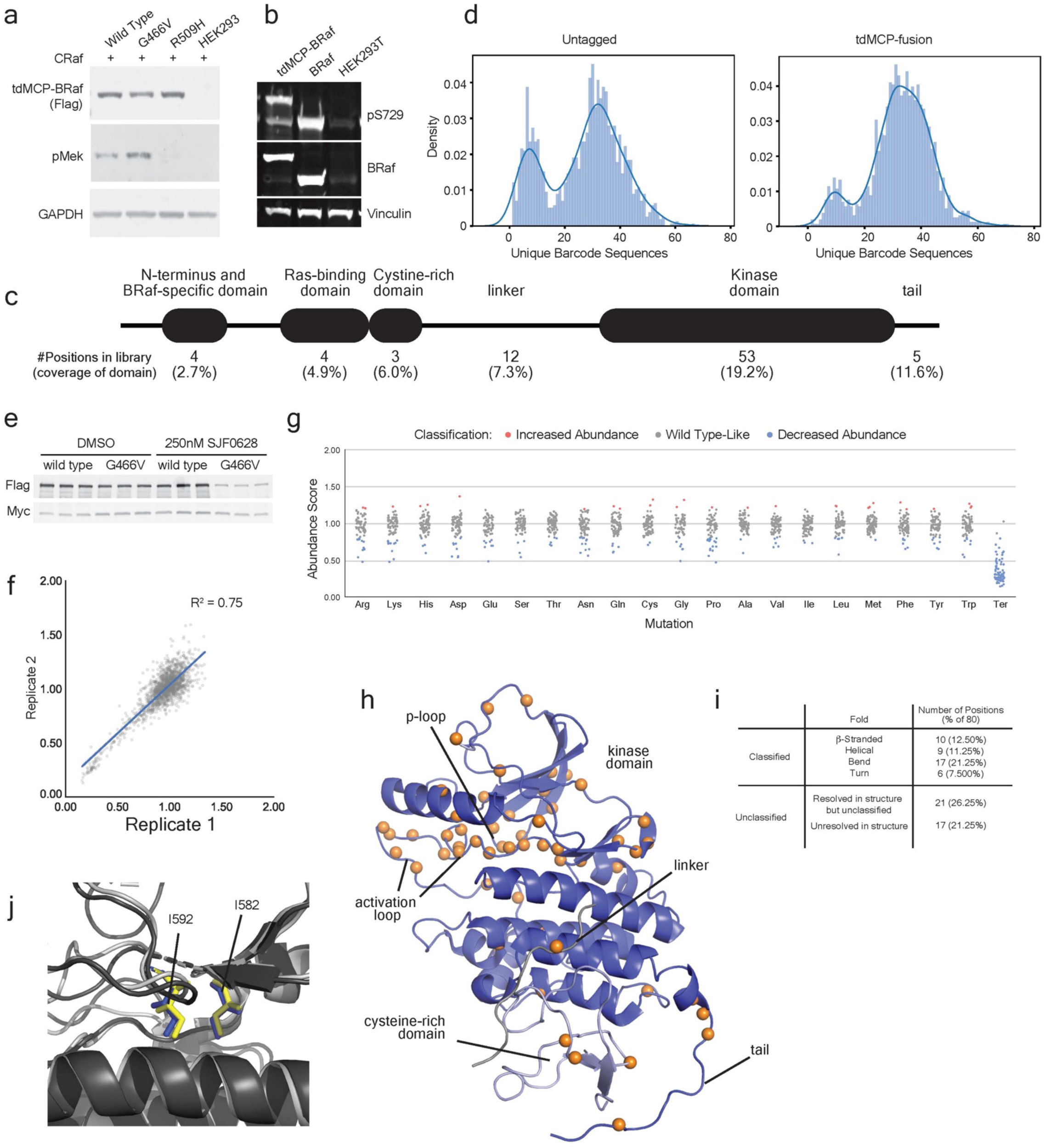
BRaf variant library construction and abundance. **a,** Western blot analysis of RPLP cells co-expressing tdMCP-BRaf variants and CRaf. **b,** Western blot analysis of HEK293T cells stably expressing Flag-tdMCP-BRaf or untagged BRaf. **c,** Domain architecture of BRaf and the number of positions in each domain included in the tdMCP- and untagged BRaf variant libraries. Positions were chosen by compiling all missense and nonsense BRaf variants appearing in the COSMIC database and The Cancer Genome Atlas and choosing positions where there are records of (1) at least 3 total variants and (b) at least 2 unique variants. **d,** Distributions of unique barcode sequences assigned to each variant in the MCP-tagged and untagged BRaf libraries. **e,** Western blots of data shown in Fig. 3d. **f,** Scatterplot showing abundance score correlations between the two independent biological replicates shown in Fig. 3e (Pearson’s R^2^ = 0.75). **g,** Abundance scores for all variants in the tdMCP-BRaf variant library, grouped by substitution. **h,** Varied positions in the untagged and tdMCP-BRaf variant libraries are shown as orange spheres on autoinhibited BRaf (PDB ID: 6NYB). **i,** DSSP (Define Secondary Structure of Proteins) structural predictions for BRaf library positions resolved in 6NYB. DSSP predictions for structures 7MFE and 5VR3 were referenced for positions not resolved in 6NYB. **j,** Alignment of BRaf’s ATP-binding pocket in the active (black, PDB 4MNE) and autoinhibited (gray, PDB 6NYB) conformations, with the two positions (I582 and I592) with the lowest average abundance scores shown as sticks.

**Extended Data Fig. 4.**
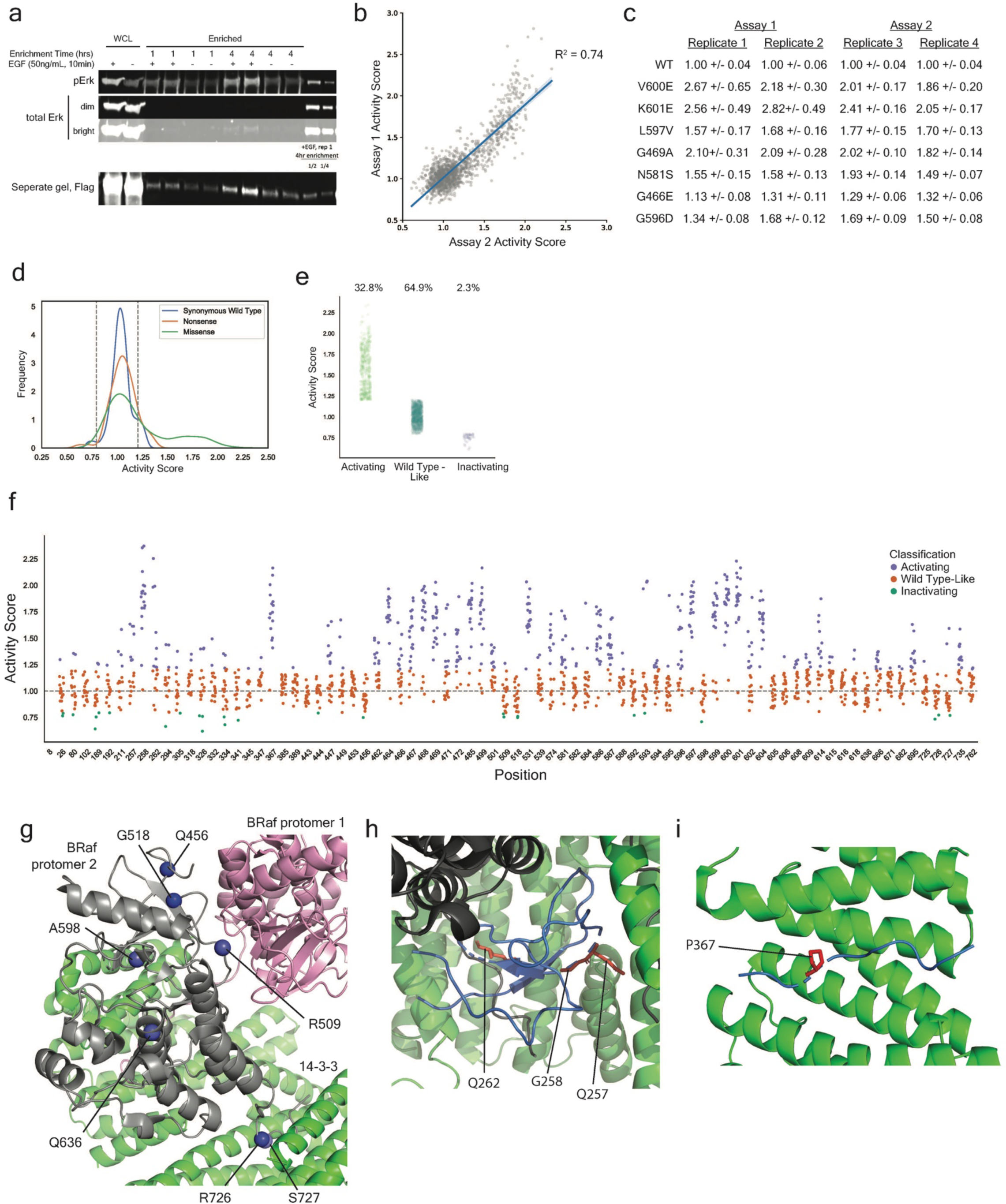
BRaf variant activity measured with overexpressed CRaf. **a,** Western blot analysis of HEK293T cells expressing the pErk reporter and treated with EGF (50 ng/mL, 10 min). Based on quantitative enrichment of pErk, it appears only a small fraction of the pErk reporter is phosphorylated under this regime. **b,** Scatterplot showing activity score correlations (Pearson’s R^2^ = 0.74) between the averages of replicates 1 and 2 with overexpressed CRaf and the averages of replicates 3 and 4 with overexpressed CRaf. Each replicate involved independent cell culturing, lysis, parallel pErk and Flag immunoprecipitations, and quantification with high-throughput sequencing. Replicates 1 and 2 underwent separate transfections with CRaf and the pErk reporter than replicates 3 and 4. **c,** Activity scores with overexpressed CRaf for previously characterized activating BRaf variants. Values shown are the mean +/- SEM of all barcodes representing a BRaf variant. **d,** LABEL-seq activity score distributions for all synonymous WT and nonsynonymous BRaf protein variants with overexpressed CRaf. The gray dashed lines indicate activity score values (>2 standard deviations of the synonymous WT score distribution higher or lower than 1, the WT score) we defined as activating and inactivating. **e,** Activity scores (with overexpressed CRaf) for BRaf variants classified as activating (32.8%), Wild Type-like (64.9%), or inactivating (2.3%). **f,** Activity scores (with overexpressed CRaf) for every BRaf variant at each position. **g,** The 10 positions with the lowest average activity scores shown as blue spheres on an active BRaf complex (PDB ID: 6Q0K). Positions 26, 189, and 443 are not resolved in the structure. **h,** The autoinhibited BRaf complex (PDB ID: 6NYB) with BRaf’s CRD shown in blue, non-CRD BRaf positions shown in black, and 14-3-3 shown in green. G258 is positioned within the CRD, which plays a role in autoinhibition, Ras binding, and membrane localization^40^. In autoinhibited BRaf, the phi-psi angles at position G258 are only energetically accessible for glycine, suggesting that substitutions at this position alter the backbone conformation and may disrupt autoinhibitory contacts, thereby leading to activation. **i,** Autoinhibited BRaf (PDB ID: 6NYB) with BRaf P367 shown as red sticks, non-P367 BRaf positions shown in blue and 14-3-3 in green. P367 is positioned in the CRD-kinase domain linker that interacts with 14-3-3 in the autoinhibited BRaf complex. Based on the known binding specificity of 14-3-3 proteins^42^, we speculate that P367 variants lead to BRaf activation by disrupting autoinhibitory 14-3-3 interactions.

**Extended Data Fig. 5.**
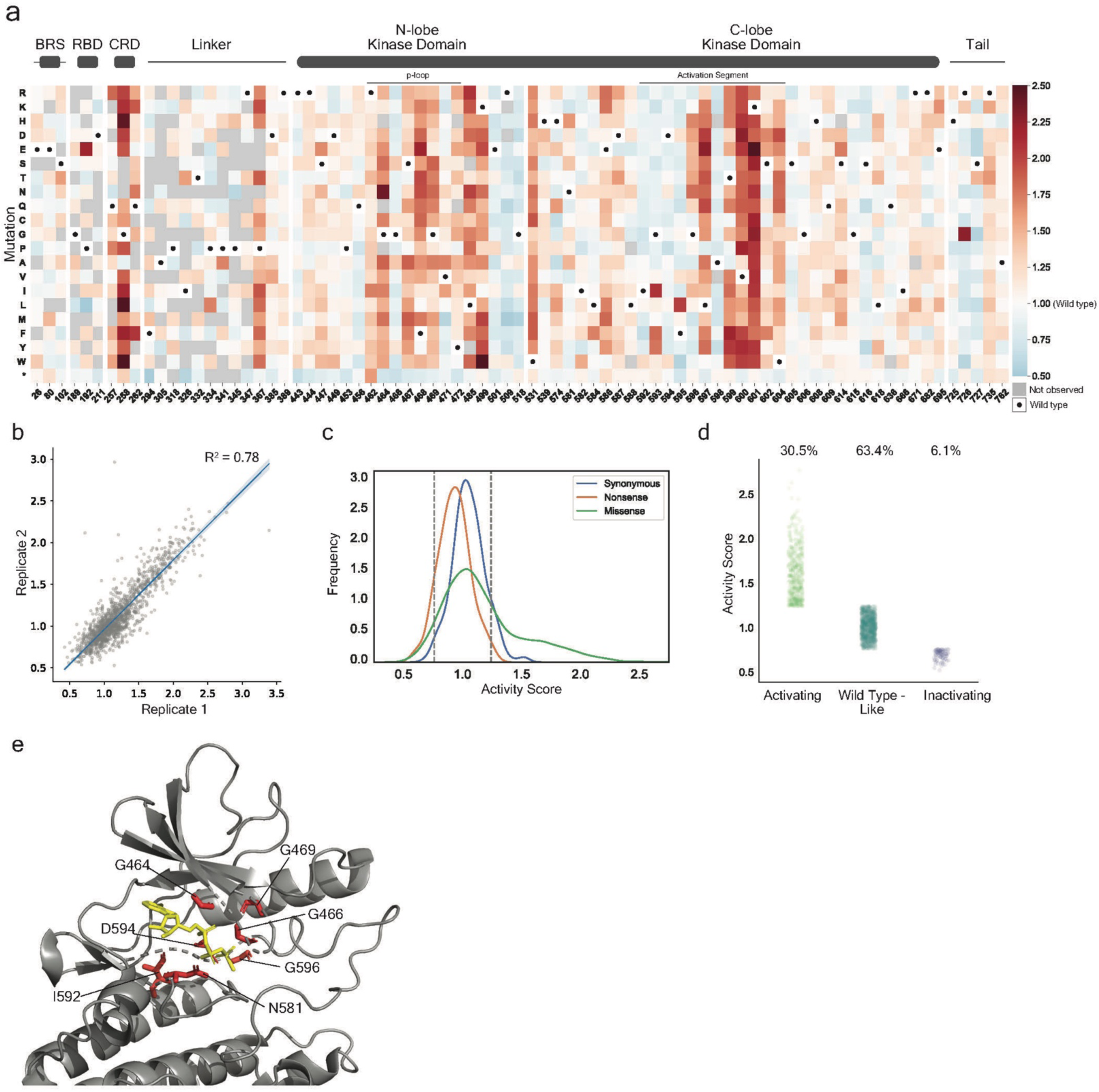
BRaf protein variant activity measured without overexpressed CRaf. **a,** Sequence-activity map for BRaf variants in the tdMCP-BRaf library measured without overexpressed CRaf. The activity scores shown are the average of two replicates, each involving independent pErk reporter expression, cell lysis, parallel Flag and pErk immunoprecipitations, and quantification by high-throughput sequencing. Black dots in the map indicate the WT amino acid and gray tiles indicate missing data. Position-averaged abundance scores are shown in the bar graph above the heatmap. **b,** Scatterplot showing activity score (without CRaf) correlations between the two independent replicates shown in **a** (Pearson’s R^2^ = 0.78). **c,** LABEL-seq activity score (without CRaf) distributions for all synonymous WT and nonsynonymous BRaf protein variants. The gray dashed lines indicate the activity score values (>2 standard deviations of the synonymous WT score distribution higher or lower than 1, the WT score) defined as increased and decreased activity. **d,** Activity scores (without CRaf) for BRaf variants classified as activating (30.5%), Wild Type-like (63.4%), or inactivating (6.1%). **e,** Positions highly conserved across the human kinome are represented as red sticks on a structure of BRaf bound to ATP-γ-S (PDB ID: 6NYB).

**Extended Data Fig. 6.**
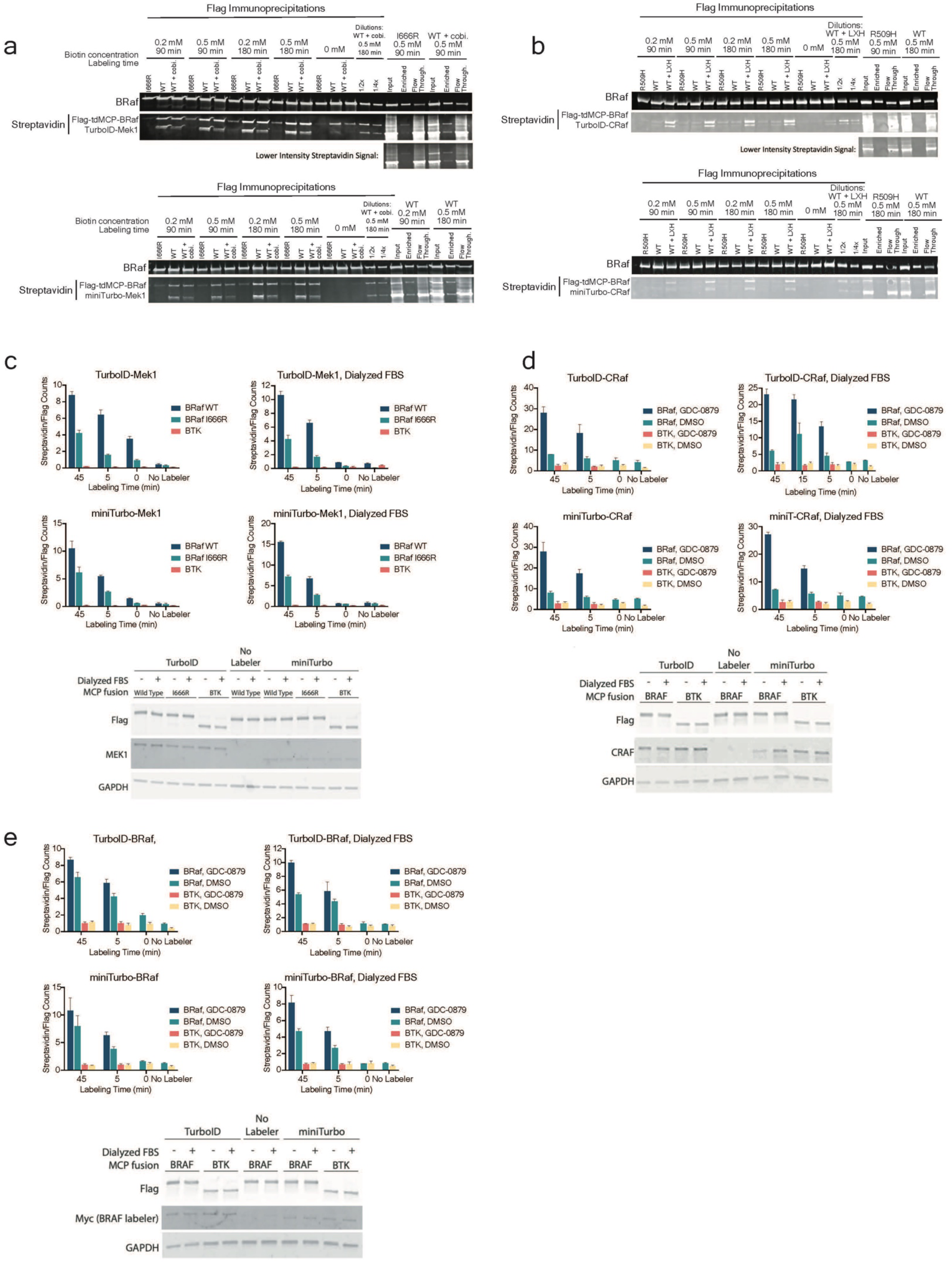
Optimization of proximity labeling conditions. **a,** Biotinylation levels of Flag-tdMCP-BRaf variants in HEK293 cells co-expressing TurboID-Mek1 (top) or miniTurbo-Mek1 (bottom), treated with 10 μM cobimetinib or DMSO for 4 h, and labeled with biotin. **b,** Biotinylation levels of Flag-tdMCP-BRaf variants in HEK293 cells co-expressing TurboID-CRaf (top) or miniTurbo-CRaf (bottom), treated with 10 μM LXH254 or DMSO for 2 h, and labeled with biotin. (**c-e**), Biotinylation levels (represented as the ratio of streptavidin and Flag barcode sequence counts) of Flag-tdMCP-BRaf variants and the background control Flag-tdMCP-BTK in HEK293T cells co-expressing TurboID-Mek1 (**c**), TurobID-CRaf (**d**), or TurboID-BRaf (**e**) and MS2-circRNAs. Cells were cultured in regular or dialyzed FBS and treated with biotin for 0, 5, or 45 min. Following biotin treatment, cells were lysed and pooled, parallel streptavidin and Flag enrichments were performed, co-enriched barcodes were sequenced by high-throughput sequencing, and the ratio of streptavidin and Flag counts for each co-enriched MS2-circRNA barcode sequence was calculated (n=3 barcodes/treatment, values shown = mean, error bars represent +/- SEM). Whole-cell lysate samples were reserved for western blots, which are shown below the bar graph (corresponds to the data shown in Fig. 5c, 5e, and 5f as well). TurboID-Mek1 (dialyzed FBS), TurboID-CRaf (dialyzed FBS), and TurboID-BRaf (dialyzed FBS) data are the same as in Fig. 5c**,e****,f** respectively.

**Extended Data Fig. 7.**
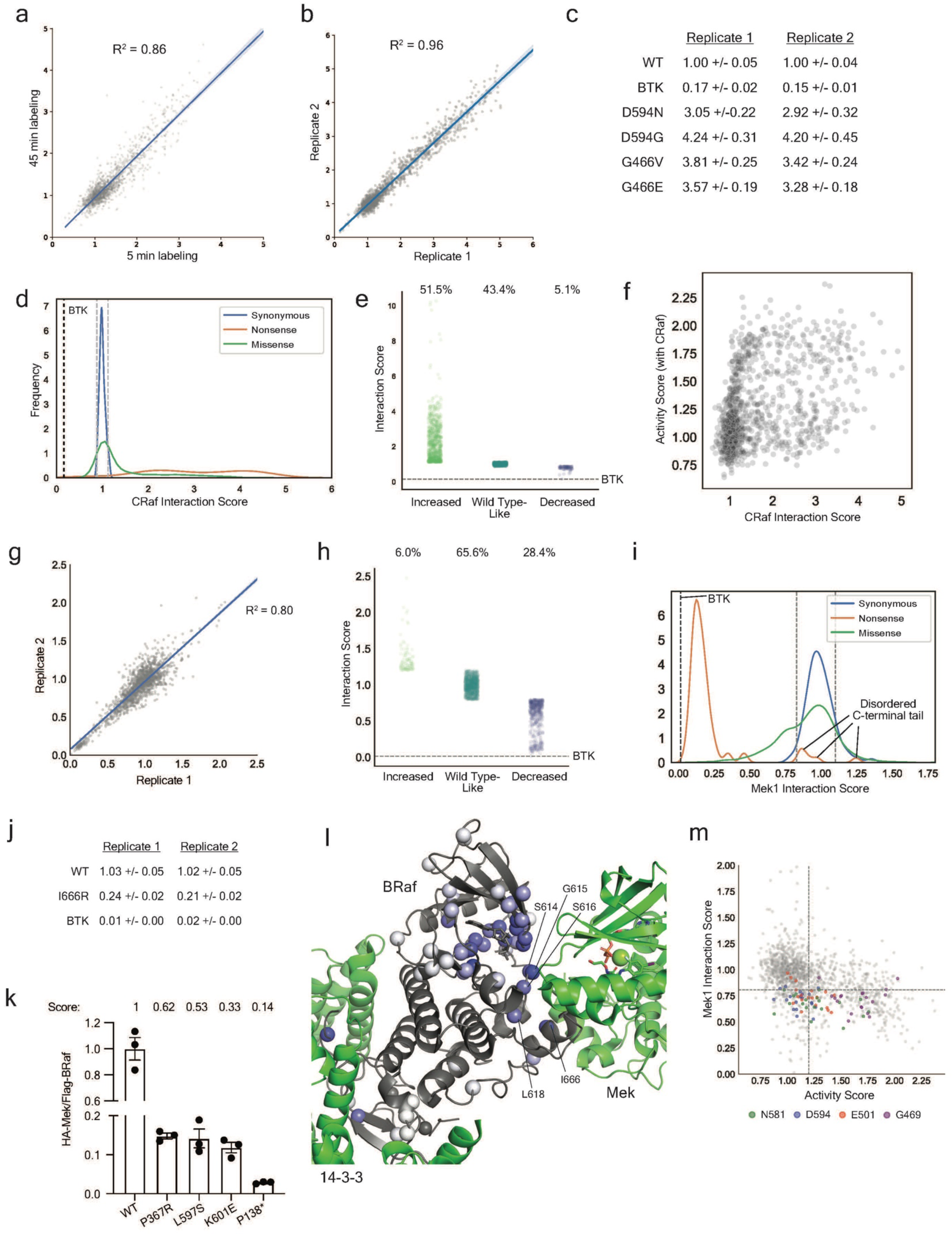
Quantifying BRaf variant interactions with CRaf and Mek1. **a,** Scatterplot showing CRaf interaction score correlations between 5 min and 45 min biotin labeling regimes (Pearson’s R^2^ = 0.86). Each score is the average of two replicates corresponding to parallel biotin labeling, streptavidin and flag enrichments, and quantification by high-throughput sequencing. **b,** Scatterplot showing CRaf interaction score correlations between the two 5 min biotin labeling replicates shown in Fig. 5h (Pearson’s R^2^ = 0.96). **c,** CRaf interaction scores shown in Fig. 5h for BRaf variants previously shown to exhibit elevated heterodimerization with CRaf. Values shown are the mean +/- SEM of all barcodes representing a BRaf variant. **d,** LABEL-seq CRaf interaction score distributions for all synonymous WT and nonsynonymous BRaf variants. The gray dashed lines indicate the CRaf interaction score values (>2 standard deviations of the synonymous WT score distribution higher or lower than 1, the WT score) we defined as increased and decreased interaction, respectively. The black dashed line is the BTK score. **e,** CRaf interaction scores for BRaf variants classified as increased CRaf interaction (51.5%), Wild Type-like (43.4%), or decreased CRaf interaction (5.1%). **f,** Scatterplot comparing activity scores (with CRaf) to CRaf interaction scores. **g,** Scatterplot showing Mek1 interaction score correlations between the two independent replicates shown in Fig. 5j (Pearson’s R^2^ = 0.80). **h,** Mek1 interaction scores for BRaf variants classified as increased Mek1 interaction (6.0%), Wild Type-Like (65.6%), or decreased Mek1 interaction (28.4%). **i,** LABEL-seq Mek1 interaction score distributions. The gray dashed lines indicate the Mek1 interaction score values (>2 standard deviations of the synonymous WT score distribution higher or lower than 1, the WT score) we defined as increased and decreased interaction, respectively. The black dashed line is the BTK score. **j,** Mek1 interaction scores for the decreased interaction control variant I666R and Flag-tdMCP-BTK. Values shown are the mean +/- SEM of all barcodes representing BRaf I666R or BTK.. **k,** Co-immunoprecipitations of HA-Mek1 and Flag-BRaf variants (without tdMCP). LABEL-seq Mek1 interaction scores are shown above the plot (n=3, value shown = mean, error bars = SEM). **l,** The number of variants at each position that were classified as decreased Mek1 interaction in Fig. 5j projected onto a structure of the autoinhibited BRaf complex (PDB ID: 6NYB). White = 0 decreased interaction variants, darkest blue = 19 decreased interaction variants. **m,** LABEL-seq Mek1 interaction and activity (with CRaf) scores. Variants at select positions within the ATP-binding pocket are highlighted. Notably, these BRaf variants possessed a range of activity scores but similarly decreased Mek1 interaction scores, indicating that differences in Mek1 interactions cannot solely be attributed to the formation of pMek, which has a lower affinity for BRaf^49^. Scores to the right of the vertical dashed line are classified as activating and scores below the horizontal dashed line are classified as decreased Mek1 interaction.

**Extended Data Fig. 8.**
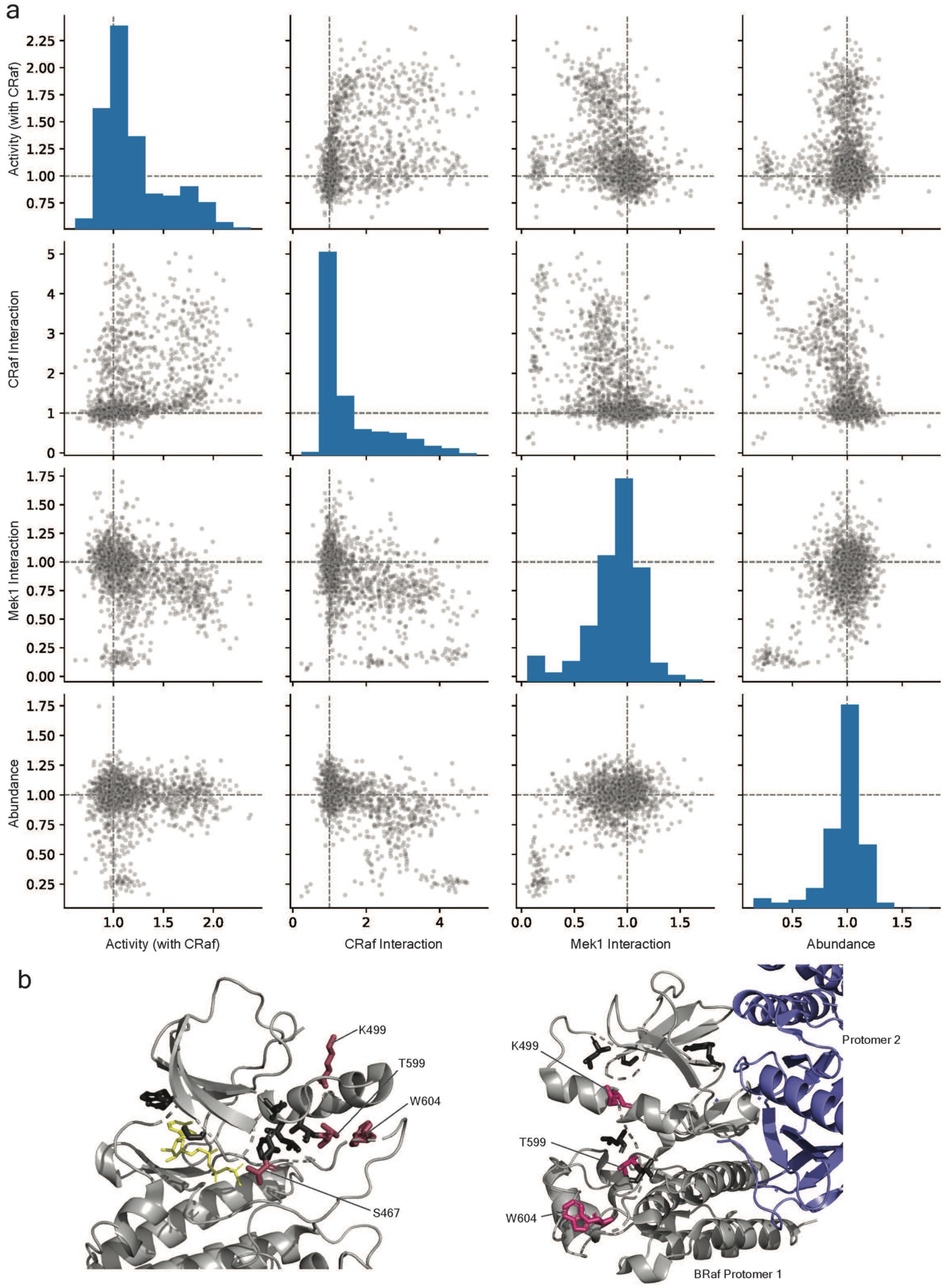
Pairwise comparisons of protein variant scores. **a,** All pairwise comparisons of activity scores (with CRaf), CRaf interaction scores, Mek interaction scores, and abundance scores. The histograms are the relative frequency of each score in the library. **b,** Cluster 1 (black sticks) and cluster 2 (maroon sticks) positions shown on autoinhibited BRaf (left, PDB ID: 6NYB) and the active BRaf dimer (right, PDB ID: 4MNE).

**Extended Data Fig. 9.**
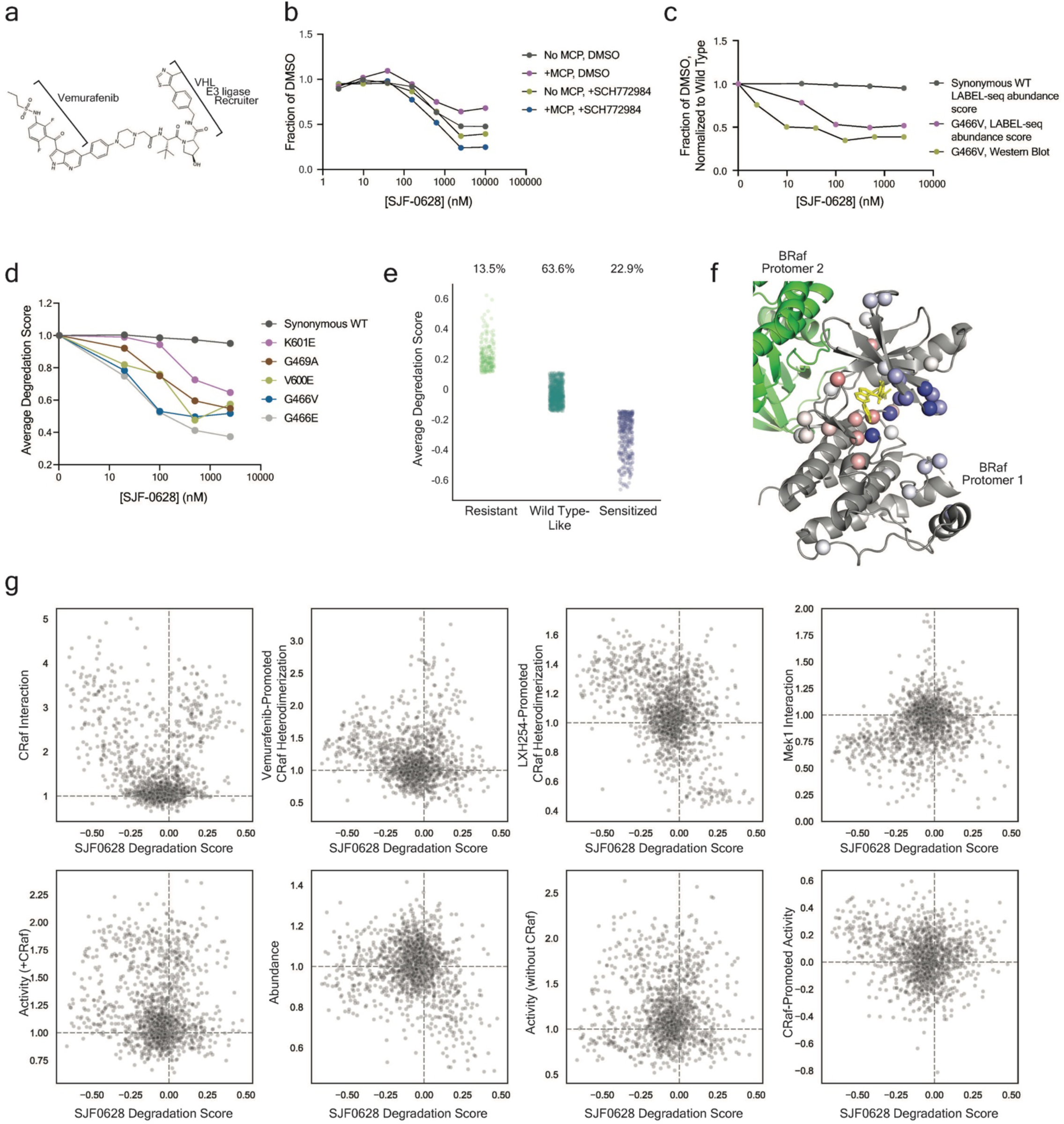
SJF0628-mediated degradation of BRaf protein variants. **a,** Structure of SJF0628. **b,** SJF0628-induced degradation of BRaf WT expressed with and without a tdMCP fusion in HEK293T cells. **c,** SJF0628-induced degradation of tdMCP-BRaf synonymous WT and G466V variants measured using western blotting and the LABEL-seq abundance assay. **d,** Average LABEL-seq SJF0628 degradation scores for five BRaf variants previously shown to be sensitive to SJF028 (n=2). **e,** Degradation scores for BRaf variants classified as resistant (13.5%), Wild Type-like (63.6%), or sensitized (22.9%). Variants with degradation scores more than two standard deviations of the synonymous distribution above or below WT were classified as resistant or sensitized, respectively. **f,** SJF0628 degradation scores at 2.5 μM SJF0628 at each position in the tdMCP-BRaf variant library (represented as spheres) shown on BRaf’s kinase domain bound to Vemurafenib (PDB ID: 3OG7). The average degradation score at a position is represented by the shade of red or blue: darkest red = 0.18, darkest blue = - 0.51. **g,** Scatterplots showing relationship between SJF0628 degradation score and other LABEL-seq biochemical measurements. Data shown for missense variants only. Gray dashed lines correspond to the WT score.

**Extended Data Fig. 10.**
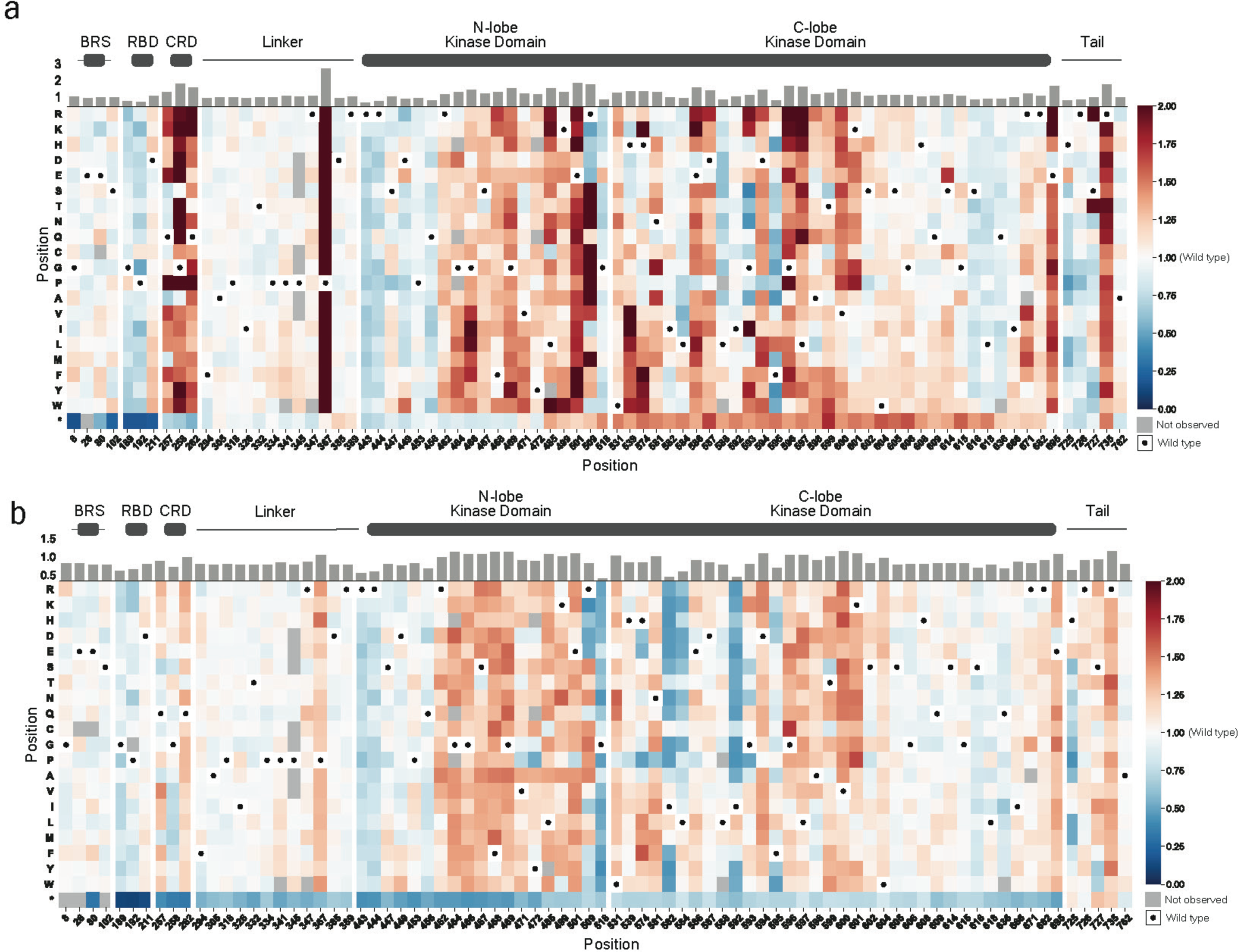
Small molecule-promoted BRaf:CRaf heterodimerization. **a-b,** Sequence-CRaf interaction maps for variants in the tdMCP-BRaf library co-expressed with TurboID-CRaf, treated with Vemurafenib (a) or LXH254 (b), and labeled with biotin. The interaction scores shown are the average of two replicates, each involving an independent cell culturing, biotin labeling, parallel streptavidin and Flag enrichments, and quantification with high-throughput sequencing. Black dots in the map indicate the WT amino acid and gray tiles indicate missing data. Position-averaged CRaf interaction scores are shown in the bar graph above the heatmaps.

## Methods

### Cloning

All plasmids, except those encoding libraries, were constructed using gibson assembly and standard protocols then validated through Sanger or whole-plasmid sequencing.

### Expression of MS2-circRNA architectures

HEK293T cells were seeded to 80% confluent in 12-well plates and transfected with a mix of 0.7 μg of plasmid encoding MS2-circRNA barcode + 0.1 μg pMAX-GFP (to estimate transfection efficiency) + 2.4 μL PEI. 48 hrs after transfection, media was removed, cells were resuspended directly in TRIzol (Invitrogen, Cat #15596026), and total RNA was extracted according to the manufacturer’s instructions. Total RNA concentrations were normalized using a NanoDrop 2000 (Thermo Scientific).

### Pulldowns with gel-based visualization of MS2-circRNA

To prepare dasatinib-conjugated sepharose, 1.25 mL of NHS-activated sepharose slurry (Cytiva cat #17090601) was washed twice with 50% DMF/50% EtOH then incubated with 64 μL 50mM dasatinib-amine + 240 μL 1M EDC + 1.6 mL 50% DMF/50% EtOH overnight at room temperature rotating end-over-end. After coupling, beads were washed three times with 50% DMF/50% EtOH then capped with ethanolamine by incubating with 544 μL 1M ethanolamine + 240 μL 1M EDC + 1056 μL 50% DMF/50% EtOH overnight at room temperature rotating end-over-end. Capped beads were washed three times with 50% DMF/50% EtOH, two times with 0.5M NaCl, then one time with 20% ethanol in water. Washed beads were resuspended in 20% ethanol to make a 50% bead/50% solvent slurry and stored at 4°C.

For pulldowns, HEK293T’s were seeded to 80% confluent in 12-well plates and transfected with a mix of 0.75 μg tdMCP-protein fusion plasmid + 0.75 μg MS2-circRNA plasmid + 0.15 μg pMAX-GFP (to estimate transfection efficiency) + 5 μL PEI. 48 hrs after transfection, cells were resuspended in 160 μL lysis buffer (50mM Tris pH=7.6, 150mM NaCl, 10mM NaF, 4mM MgCl_2_, 5% glycerol, Halt protease inhibitor tablet (1 tablet/50mL, Fisher cat# A32965), 2mM PMSF, RiboLock RNase Inhibitor (4 μL/mL, Fisher cat#FEREO0382)) and lysates were cleared at 17,000g for 10 min at 4°C. Prior to enrichment, 40 μL of supernatant was incubated with 15 μM free dasatinib (1% DMSO) or 1% DMSO for 30 min at 4°C. To enrich, 10 μL of dasatinib-conjugated sepharose (pre-washed with lysis buffer) was added to each sample and incubated for 2.5 h at 4°C, rotating end-over-end. Protein-bound beads were washed three times with 100 μL of lysis buffer. For western blot analysis, beads were resuspended in 3xSDS. For analysis of RNA enrichment, beads were resuspended in 100 μL TRIzol (Invitrogen cat# 15596026). All samples were stored at -20°C. RNA was extracted and purified using RNA cleanup columns (IBI scientific TRI-Isolate RNA Pure Kit, VWR cat# 76006-526), eluted into 25 μL of water and immediately visualized by ge (see “Gel-based visualization of RNA”)

### Gel-based visualization of MS2-circRNA

To visualize broccoli by gel, total RNA containing MS2-circRNA (1 μg for scaffold test expression, 7 μL for pulldown samples) was mixed 1:1 with 2x TBE-urea sample buffer (Invitrogen, Fisher cat #LC6876), heated to 75°C for 5 min, then briefly cooled on ice. Samples were loaded onto precast 6% TBE (Invitrogen, Fisher cat #EC62655BOX) or 10% TBE-Urea (Invitrogen, Fisher cat #EC68755BOX) gels and run at 270V in TBE buffer to completion. Gels were washed three times for 5 min in milliQ water then stained for 20 min at room temperature in DFHBI buffer (10 μM DFHBI-1T, 40mM HEPES, pH 7.4, 100mM KCl, and 1mM MgCl2), both steps on an orbital shaker. Broccoli was imaged on a Typhoon FLA 9000 using preset settings for Alexa Fluor 488 (495nm excitation and 519nm emission).

### Generating the RNA-Protein Landing Pad (RPLP)

To stably generate a high-efficiency Bxb1 landing pad containing Pol II- and Pol III-driven promoters, Flp-In™ T-REx™ 293 cells (Fisher cat#R78007) were plated into a 6-well dish and transfected with 1 μg of landing pad plasmid containing an FRT site, 1 μg Flp recombinase expression plasmid (pOG44), 6 μL PEI, and 100 μL serum-free DMEM. 48 hrs after transfection, cells were re-seeded to ∼20% confluent. After an additional 24 hrs, media was supplemented with 200 μg/mL hygromycin (Mirus Bio cat#MIR 5930) to initiate selection. Cells were expanded and maintained in 200 μg/mL hygromycin until >95% of cells were GFP positive indicating near-complete selection, and stocks were cryo-preserved until needed.

### Cloning single variant barcode libraries

To generate a library of barcodes, attB plasmid containing genes for tdMCP-protein fusions and a MS2-circRNA barcode were digested overnight at 37°C to remove existing barcode sequence (2 μg plasmid, 5 μL 10x CutSmart buffer, 2 μL BsrGI-HF (NEB cat#R3575S), nuclease-free water to 50uL). The next day, fragments were dephosphorylated by adding 5 μL antarctic phosphatase buffer and 2 μL enzyme (NEB cat#M0289S) and incubating for 1 h at 37°C. Fragments were then separated by gel electrophoresis and the band corresponding to digested vector was extracted using a Zymo kit (Zymo cat#D4002) and eluted into 20 μL nuclease-free water.

To generate a double-stranded barcode fragment (dsBC), 50pmol of the single stranded barcode oligo (JJS29), 50pmol of the second strand synthesis primer (ssSynth primer, JJS30), 5 μL NEB2.1 10x buffer, and nuclease-free water to a total volume of 48 μL were combined. Oligos were annealed by heating to 94°C for 5 min, ramping 0.1°C/s to 71°C (Tm) and holding for 5 min, then ramping 0.1°C/s to 37°C. ssSynth primer was extended by adding 1 μL 10mM dNTP’s (Fisher cat#FERR0192) and 1 μL Klenow polymerase (NEB cat#M0210S) then incubating at 37°C for 1hr. The resulting dsBC was cleaned using a Zymo column (Zymo cat#D4014) and eluted into 12 μL nuclease-free water.

dsBC was cloned into the digested vector by combining 200ng vector (∼0.035pmol, 8uL), 10ng dsBC (∼0.175pmol, 2uL) and 10 μL 2x NEBuilder® HiFi DNA Assembly Master Mix (NEB cat#E2621L) and incubating at 50°C for 1hr. A separate vector-only reaction was run in parallel using 200ng vector, 10 μL NEBuilder Master Mix, and nuclease-free water to 20 μL (without barcode) to estimate vector-only background transformants. After incubation, reactions were cleaned (Zymo cat#D4014) and eluted into 10 μL nuclease-free water.

Cleaned reactions were transformed into NEB® 5-alpha Competent *E. coli* (NEB cat#C2987H) following the manufacturer’s instructions. Following outgrowth, transformed cells were added to 50 mL LB broth supplemented with 100 μg/mL carbenicillin. The library was bottlenecked to the desired number of barcodes by making serial dilutions of the culture in LB+carb. 200 μL of each dilution was plated on LB+carb plates to enumerate transformants. Liquid cultures were grown at 37°C overnight to saturation and DNA was extracted using QIAGEN® Plasmid Plus Midi Kit (Qiagen cat#12945)

### Generating stable cell lines

Stable cell lines expressing protein and RNA barcodes were generated in the HEK293 RPLP. Stable lines not expressing circRNA barcodes were generated in the HEK293T lenti landing pad (LLP-iCasp9-Blast^24^). The following protocol applies to both cell lines.

Cells were maintained in high glucose DMEM (Fisher cat #11-965-118) supplemented with 10% FBS and 2 μg/mL doxycycline (DOX, Sigma cat#D9891, 0.01% DMSO final concentration). Genes of interest were cloned into Bxb1 attB-containing plasmids using standard Gibson assembly cloning techniques (NEB cat #E2621L). To stably recombine the attB plasmids under control of the landing pad promoters, cells were plated the day before transfection in 6-well dishes to 40% confluence using DOX-free media. The following day, cells were transfected with a mixture of 1.7 μg attB plasmid + 0.2 μg pCAG-NLS-Bxb1 (Addgene#51271) + 0.2 μg pMAX-GFP + 7.8 μL Fugene 6 (for LLP lines, Promega cat#E2691) or Fugene HD (for RPLP lines, Promega cat#E2311) + 100 μL serum-free DMEM (Fisher cat #11-965-118). 48-hours after transfection, cells were expanded to 10cm dishes in media containing 2 μg/mL DOX. 24-48 hrs after DOX induction, media was supplemented with 10nM AP1903 (Fisher cat#50-115-1810) to select for successful recombinants. 12-24 hrs after addition of AP1903, media was exchanged for AP1903-free media (+2 μg/mL DOX), the resulting stables were expanded in +DOX media, and stocks frozen until required.

### Barcode sequence missassociation experiment

Two barcode libraries were cloned as described in the ‘Generating single protein variant barcode libraries’ section, each containing 20,000-30,000 unique barcode sequences and either 3xFlag- or 2xMyc-tdMCP-BRaf wild type. Plasmid libraries were individually recombined into the RPLP as described in the ‘Generating stable lines’ section.

The day before enrichment, equal numbers of cells from both libraries were combined and plated into 15 cm dishes in +DOX media. On the day of enrichment, cells were trypsinized and all plates pooled. The pool aliquots were then divided into 5 enrichment samples: 2 no-dummy samples (30×10^6^ cells each), 2 +dummy samples (10×10^6^ cells each), and 1 input sample (10×10^6^ cells each). Each aliquot was washed once with DPBS, then fully resuspended in 1,650 μL (no dummy samples) or 550 μL (+dummy and input) modRIPA without detergent (i.e. resuspension buffer: 50mM Tris pH=7.6, 150mM NaCl, 10mM NaF, 4mM MgCl_2_, 5% glycerol, Halt protease inhibitor tablet (1 tablet/50mL, Fisher cat# A32965), 2mM PMSF, RiboLock RNase Inhibitor (4 μL/mL, Fisher cat#FEREO0382)). For +dummy samples, the resuspension buffer also contained 400ng/ μL total RNA previously extracted from HEK293T cells which were transiently transfected with dummy MS2-circRNA (an MS2-circRNA architecture which contains the MS2 hairpin but the reverse transcription priming sites). Immediately after resuspending cells in resuspension buffer, cells were lysed by adding an equal volume of modRIPA containing 0.1% NP40. Addition of buffer with detergent marked the beginning of the incubation time. Lysates were cleared by centrifugation at 17,000g for 10 min at 4°C. Each enrichment sample was split into two aliquots (1.5 mL ea. for no dummy, 0.5 mL ea. for +dummy) and transferred to low-bind eppendorf tubes (Fisher cat#13-698-792). To enrich tdMCP- protein fusions, 150 μL (no dummy) or 50 μL (+dummy) of anti-flag (Sigma cat#M8823) or anti-myc (Fisher cat#PI88842) beads were added to the lysates and rotated end-over-end at 4°C for the indicated time. 500 μL samples of bead suspension were collected at each timepoint and washed 3 times with 500 μL modRIPA (including 0.1% NP40). After washing, all buffer was removed before resuspending beads in 75 μL nuclease-free water and 300 μL TRIzol, then stored at -20. The input sample was resuspended in 1.5 mL TRIzol and stored at -20. RNA was extracted using IBI scientific TRI-Isolate RNA Pure Kit, concentrated using Zymo RNA Clean & Concentrator (Zymo Research Cat #R1015), eluted into 10 μL nuclease-free water, and stored at -80°C.

### Variant abundance assay, proof of concept experiment

HEK293T LLP cells stably expressing 3xFlag-tdMCP-BRaf wild type or G466V were transfected in 24-well plates, each well receiving 300 ng pcDNA5 2xMyc-tdMCP-SNAP + 200 ng of an equal mix of 4 plasmids, each containing a unique circRNA barcode + 50 ng pMAX GFP + 50 μL DMEM + 1.65 μL PEI. 6 transfections per variant encompassing a total of 48 unique barcodes were performed. 24 hrs after transfection cells were treated with SJF0628 or DMSO for an additional 24 hrs. After drug treatment, cells were lysed in 120 μL modRIPA (0.1% NP40, 50mM Tris pH=7.6, 150mM NaCl, 10mM NaF, 4mM MgCl_2_, 5% glycerol, Halt protease inhibitor tablet (1 tablet/50mL, Fisher cat# A32965), 2mM PMSF, RiboLock RNase Inhibitor (4uL/mL, Fisher cat#FEREO0382)) and lysate cleared by centrifugation at 17,000g for 10 min at 4°C. Samples were taken for western blot analysis before pooling 40 μL lysate from all 12 wells. Pooled lysate was subsequently split into 2×200 μL aliquots and protein was enriched by adding 20 μL anti-flag (Sigma cat#M8823) or anti-myc (Fisher cat#PI88842) beads and rotating end-over-end at 4°C for 1 hr. Beads were washed 3 times with modRIPA, resuspended in 50 μL nuclease-free water and 200 μL TRIzol, then stored at -20°C. Enriched RNA was extracted and sequenced as described in the ‘Preparation of RNA for Genewiz Amplicon-EZ sequencing’ section.

Sequencing ratios were computed by dividing the sequencing counts in the flag enrichment by counts in the Myc enrichment then normalizing to the mean flag/myc ratio for wild type cells treated with DMSO. The mean ratio of all barcodes in a well was reported. Similarly, for western blot ratios the intensity of the flag signal was divided by the intensity of the Myc signal. Ratios were normalized to the average ratio of wild type cells treated with DMSO. The three data points correspond to the three transfections.

### Variant activity assay, proof of concept experiment

HEK293T LLP cells stably expressing untagged BRaf wild type or V600E were transfected in 12-well plates with 0.5 μg 3xFlag-tdMCP-Erk2 + 0.4 μg of an equal mix of 4 plasmids, each containing a unique circRNA barcode + 0.1 μg pMAX-GFP + 50 μL DMEM + 3 μL PEI. Transfections of each variant were performed in triplicate, encompassing a total of 24 unique barcodes. 48 hrs after transfection cells were lysed in 150 μL modRIPA (0.1% NP40, 50mM Tris pH=7.6, 150mM NaCl, 10mM NaF, 4mM MgCl_2_, 5% glycerol, Halt protease inhibitor tablet (1 tablet/50mL, Fisher cat# A32965), 2mM PMSF, RiboLock RNase Inhibitor (4 μL/mL, Fisher cat#FEREO0382), 100 μL each phosphatase inhibitor cocktails 2 and 3 (Sigma Cat #P5726 and P0044)) and lysate cleared by centrifugation at 17,000g for 10 min at 4°C. 30 min after lysis all samples were pooled. The pooled lysate was divided into two 350 μL aliquots and protein was enriched by adding 35 μL anti-flag (Sigma cat#M8823) or anti-pErk beads (see ‘Basal Activity Scan’ for instructions on generating pErk beads), rotating end-over-end at 4°C for 2 hr. Beads were washed 3 times with modRIPA, resuspended in 50 μL nuclease-free water and 200 μL TRIzol then stored at -20°C. Enriched RNA was extracted and sequenced as described in the ‘Preparation of RNA for Genewiz Amplicon-EZ sequencing’ section.

pErk/Flag ratios were computed by dividing the sequencing counts in the flag enrichment by counts in the pErk enrichment. The mean ratio of all barcodes in a well was reported.

### Proximity labeling time course

Prior to the experiment, MS2-circRNAs 1b comprising 48 unique barcode sequences were expressed individually in HEK293T’s. 48 hrs after transfection, cells were collected and total RNA was extracted using TRIzol (Invitrogen, Cat #15596026) and isopropanol precipitation following the manufacturer’s instructions. The extracted total RNA was diluted to ∼1μg/μL in nuclease-free water, aliquoted, and stored at -80.

HEK293T LLP cells stably expressing tdMCP fusions (BRaf wild type, BRaf I666R, or BTK) were seeded into plates using DMEM supplemented with regular or dialyzed FBS. Cells were carried through the entire experiment in the same type of FBS (regular or dialyzed). After seeding, cells were transfected with TurboID- or miniTurbo-fusions (Mek1, CRaf, or BRaf). Transfection mixes per 1×10^6^ cells included 1.8 μg pCDNA5 TurboID/miniTurbo plasmid + 0.2 μg pMAX GFP to estimate transfection efficiency + 6 μL Turbofectin (Origene cat# TF81001)+ 100 μL DMEM. 24 hrs after transfection, transfected cells were re-seeded to ∼80% confluency into 96-well poly-D-lysine coated plates. ∼24 hrs after moving to 96-well plates, cells were treated with DMSO or 10 μM GDC-0879. After 2 h pre-treatment with drug, biotin treatment was initiated by supplementing media with biotin to a final concentration of 150 μM biotin and 10 μM GDC-0879. Subsequent biotin labelings were initiated to achieve 5, 15, or 45 min labeling. Labeling was quenched for all samples simultaneously by removing media, washing 3x with warm DPBS, and immediately adding 40 μL cold modRIPA (50mM Tris pH=7.6, 150mM NaCl, 10mM NaF, 4mM MgCl_2_, 5% glycerol, Halt protease inhibitor tablet (1 tablet/50mL, Fisher cat# A32965), 2mM PMSF, RiboLock RNase Inhibitor (4uL/mL, Fisher cat#FEREO0382), 0.1% NP40) to each well. Cells were incubated at 4°C for 20 min on an orbital shaker to ensure cells were fully lysed. During the 4°C incubation, total RNA stocks containing MS2-circRNA’s were thawed and diluted 1/75 in cold modRIPA. 10 μL of the diluted RNA was added to each lysate-containing well, with each well receiving a unique barcode sequence. Lysates were incubated with RNA at 4°C for 2 h on an orbital shaker to equilibrate MS2-circRNA-tdMCP binding. After incubation, 50 μL cold modRIPA supplemented with 1mg/mL BSA was spiked into each well and 80 μL from each well were pooled. Lysates were cleared by centrifugation at 17,000g for 10 min at 4°C. 1.9 mL of supernatant was combined with 95 μL anti-flag beads (sigma) or streptavidin conjugated-agarose (Fisher cat# PI20361) and rotated end-over-end at 4°C for 1hr. After incubation, the beads were washed 3×1 mL cold modRIPA then resuspended in 75 μL water and 300 μL TRIzol and stored at -20°C. MS2-circRNA was extracted and sequenced as described in ‘Preparation of RNA for Genewiz Amplicon-EZ sequencing’.

### Preparation of RNA for Genewiz Amplicon-EZ sequencing

Enriched RNA was extracted from TRIzol using IBI scientific TRI-Isolate RNA Pure Kit, eluted into 25 μL nuclease-free water and stored at -80°C. SuperScript™ IV Reverse Transcriptase (Invitrogen Cat#18090010) was used to generate cDNA from circRNA barcodes, following the manufacturer’s instructions for gene-specific priming. Total reaction volume was 5 μL including 1 μL of extracted RNA and JJS1 as the primer.

To generate amplicons, 5 rounds of PCR were run using Q5® Hot Start High-Fidelity 2X Master Mix (NEB cat #M0494) following the manufacturer’s instructions with a 63°C annealing temperature and 30s extension. A 25 μL reaction scale was used including 12.5 μL Q5® Master Mix, 1.25 μL each of JJS2 & JJS3 (10 μM stocks), 2.5 μL cDNA reaction, 7.5μL nuclease-free water, and 0.75 μL DMSO. PCR reactions were cleaned with Sera-Mag Select beads (Cytiva #29343052) using a 2.5:1 ratio of beads:reaction and eluted using 25 μL nuclease-free water.

A second round of PCR was run on the eluted amplicons using 12.5 μL Q5® Master Mix, 1.25 μL each JJS4 & JJS5 (10 μM stocks), 5 μL eluted amplicon, 5 μL nuclease-free water, 1x SYBR gold (Invitrogen Cat#S11494), and 2% DMSO. Reaction progress was monitored using a Bio-Rad CFX Connect Real-Time PCR Detection System and halted in the linear phase of amplification. Reactions were equally pooled based on their final RFU values and amplicons of the correct size were selected using Sera-Mag Select beads with a 0.65:1 ratio of beads:reaction. Amplicons were eluted into nuclease free water and submitted to Genewiz using their Amplicon-EZ service for sequencing.

### Western blotting of individual variants

**(applies to library score vs quantification by western plots)**

HEK293T LLP cells stably expressing flag-tagged BRaf variants of interest were lysed in cold modRIPA (50mM Tris pH=7.6, 150mM NaCl, 10mM NaF, 4mM MgCl_2_, 5% glycerol, Halt protease inhibitor tablet (1 tablet/50mL, Fisher cat# A32965), 2mM PMSF, 0.1% NP40, 1% each phosphatase inhibitor cocktails 2 and 3 (Sigma Cat #P5726 and P0044). Lysates were cleared at 17,000g, 4°C for 10 min and total protein concentrations were measured then normalized using Pierce™ 660nm Protein Assay Reagent (Fisher cat#PI22660). Samples were mixed 2:1 with 3xSDS loading buffer (188 mM Tris-Cl (pH 6.8), 3% SDS, 30% glycerol, 0.01% bromophenol blue, 15% β-mercaptoethanol) and heated to 95°C for 5 min. 3-10 μg total protein was loaded and run on AnykD™ Criterion™ TGX™ Precast acrylamide protein gels (Biorad cat#5671125) then transferred to nitrocellulose (Biorad cat#1704159). Nitrocellulose was incubated at room temp for one hour in 5% dry milk in TBST then treated overnight with primary antibody diluted in 5% dry milk in TBST at 4°C. The next day, nitrocellulose was washed 3 times for 5 min in TBST then incubated for at least 1 h with secondary antibody (Licor cat#926-68070 or 926-32211) diluted 1:10,000 in 5% dry milk in TBST. Westerns were scanned on a LI-COR Odyssey® and quantified using Image Studio software.

pErk was probed using CST#4370L, total Erk CST#9107S, BRaf CST#14814S, Flag Sigma #F1804, GAPDH Santa Cruz Biotechnology #365062, streptavidin fluorophore LI-COR #926-32230, pMek CST cat#9154S, HA CST#3724, BRaf pS729 Abcam #ab124794, Vinculin CST#13901.

### Mek1 co-immunoprecipitations

HEK293T LLP cells stably expressing flag-tagged BRaf variants (wild type, P367R, K601E, L485S, P318*) were transfected in 6-well plates in triplicate with 1.8 μg pcDNA5 HA-Mek1 + 0.2 μg pMAX GFP + 6 μL PEI + 100 μL DMEM. 48 hrs after transfection, cells were lysed using 200 μL high-glycerol modRIPA (0.1% NP40, 50mM Tris pH=7.6, 150mM NaCl, 10mM NaF, 4mM MgCl_2_, 10% glycerol, Halt protease inhibitor tablet (1 tablet/10mL, Fisher cat# A32955), phosphatase inhibitor cocktails 2 and 3 (100uL/10mL, Sigma Cat #P5726 and P0044)) and lysates were cleared by centrifugation at 17,000g for 10 min at 4°C. To enrich Mek-Raf complexes, 150 μL of supernatant was combined with 12 μL anti-flag beads (50% slurry, washed, (Sigma cat#M8823)) and rotated end-over-end at 4°C for 1 h then quickly washed three times with 100 μL modRIPA. Enriched complexes were eluted from the bead by resuspending in 30 μL 3xSDS and heating to 95°C for 5 min.

### Cloning tdMCP-BRaf variant library

A site saturation mutagenesis library of BRaf consisting of variants at 79 chosen residues was purchased from Twist Bioscience. The double stranded DNA pool contained the entire 3xFlag-tdMCP-BRaf open reading frame with flanking 5’ and 3’ vector overlaps. The gene fragment library was cloned into the linearized attB vector using Gibson assembly.

To generate a linearized vector the following was combined and incubated at 37°C overnight: 2 μg attB library vector, 1.5 μL AflII (NEB cat#R0520S), 1.5 μL NheI-HF (NEB cat#R3131S), 5 μL CutSmart 10x buffer, nuclease-free water to 50uL. The next day, 2.5 μL recombinant shrimp alkaline phosphatase (NEB cat#M0371S) was added and incubated at 37°C for 2 hrs. Digested DNA was run on a 1% agarose gel, extracted (Zymo cat#D4008), and eluted into 30 μL nuclease-free water.

The gene fragment library was cloned into the linearized vector using a 1:2 molar ratio of vector:insert by combining the following and incubating at 50°C for 1hr: 80ng linearized vector, 95ng gene fragment library, 40 μL NEBuilder® HiFi DNA Assembly Master Mix (NEB cat#E2621L), and nuclease-free water to 80uL. To estimate background transformants, a vector-only reaction was run in parallel using 80ng linearized vector, 40 μL NEBuilder Master Mix, and water to 80 μL (no gene fragment library). After incubation, reactions were cleaned (Zymo cat#D4014) and eluted into 20 μL nuclease-free water.

Cleaned reactions were transformed into NEB® 5-alpha Competent E. coli (High Efficiency) (NEB cat#C2987H) per the manufacturer’s instructions. Briefly, 4 replicate transformations were performed on the library, each using 40 μL cells and 4 μL cleaned gibson reaction and single replicate was performed in parallel to estimate vector-only transformants. Cells were incubated with DNA on ice for 30 min, heat shocked at 42°C for 30 sec, and put back on ice for 5 min. Subsequently, 760 μL SOC was added to each replicate and cells were incubated at 37°C for 1 h at 220 rpm. All library replicates were then pooled and added to 40 mL LB supplemented with 100 μg/mL carbenicillin (Fisher cat#BP2648250). From this culture, 200uL, 20uL, and 2 μL was plated onto carbenicillin-selective agar plates to estimate transformants. 200 μL vector-only was also plated onto agar to estimate background. The 40 mL liquid culture was grown overnight to saturation at 37°C and 220rpm. The following day DNA was harvested (Qiagen cat#12145).

Variant library was barcoded and bottlenecked to desired number of transformants as described in ‘Generating single protein variant barcode libraries’.

### Cloning untagged BRaf variant library

To clone a BRaf variant library without the 3xFlag-tdMCP fusion, the attB 3xFlag-tdMCP-BRaf variant library (containing a single barcode) was digested by combining the following and incubating overnight at 37°C: 2 μg plasmid + 5 μL CutSmart buffer (10x) + 2 μL AflII (NEB cat# R0520S) + 2 μL BlpI (NEB cat# R0585S) + nuclease-free water to 50 μL. The next day, 1 μL recombinant shrimp alkaline phosphatase (NEB cat#M0371S) was added to the reaction and incubated at 37°C for 1hr. The digest was run on a 1% agarose gel, the correct fragment extracted (Zymo cat#D4008), then eluted into 30 μL nuclease-free water.

To generate a double stranded bridging fragment the following was combined: 1 μg (36.1pmol) JJS6 + 870ng (36pmol) JJS7 + 5 μL NEBuffer 2 (10x) + nuclease-free water to 48uL. Oligos were annealed by denaturing at 94°C for 5 min, ramping 0.1°C/s to 65.5°C (Tm), holding at 65.5°C for 5 min, then ramping 0.1°C/s to 37°C and holding. 1 μL dNTP’s (10mM each, NEB cat#FERR0192) and 1 μL Klenow polymerase (NEB cat#M0210S) was added to the annealed oligos. Oligos were extended by incubating at 37°C for 1hr, inactivating the enzyme at 75°C for 20 min, re-annealing the DNA by ramping 0.1°C/s to 42°C, then holding at 4°C. DNA was cleaned and concentrated (Zymo cat#D4014), then eluted into 30 μL nuclease-free water.

Digested variant library was re-circularized using gibson assembly at a 1:5 molar ratio of vector:bridge as recommended by the manufacturer. Briefly, the following was combined and incubated at 50°C for 1hr: 200ng (5 μL) digested vector, 20ng (2 μL) dsBridge, 13 μL nuclease-free water, 20 μL 2x HiFi DNA Assembly master mix (NEB cat#E2621L). After incubation, reactions were cleaned (Zymo cat#D4014) and eluted into 20 μL nuclease-free water.

Cleaned gibson reactions were transformed into NEB® 5-alpha Competent E. coli (High Efficiency) (NEB cat#C2987H), transformants enumerated, and DNA harvested as described in ‘Cloning MCP-tagged gene library into attB library vector’.

Variant library was barcoded and bottlenecked to desired number of transformants as described in ‘Generating single protein variant barcode libraries’.

### Generating variant library cell lines

Barcoded libraries were stably recombined into the HEK293 RPLP cell line, maintained in high glucose DMEM (Fisher cat #11-965-118) supplemented with 10% FBS and 2 μg/mL doxycycline (Sigma cat#D9891). The day before transfection RPLP cells were plated into two 15cm dishes using DOX-free media (+10% FBS) to reach ∼85% confluent during transfection. On the day of transfection, media was replaced with dox-free DMEM (+10% FBS) supplemented with 100nM SCH772984. Transfection mixes were prepared by combining per 15cm dish: 16.5 μg attB-plasmid library + 3 μg pCAG-NLS-Bxb1 + 2 μg pMAX-GFP + 2150 μL serum-free DMEM + 80 μL FugeneHD (Promega cat# E2311), then incubated at room temperature for 15 min and added dropwise to cells. 48 hrs after transfection cells were expanded and maintained in DMEM (+10% FBS) supplemented with 2 μg/mL doxycycline and 100nM SCH772984 moving forward. 4 days after transfection cells were further expanded and a sample analyzed by flow cytometry to quantify recombination efficiency via the mCherry reporter. 6 days after transfection, selection was initiated by supplementing media with 10nM AP1903. After 6 hrs, media was replaced with fresh +AP1903 media. 24 hrs after initiating selection, cells were further expanded in media supplemented with 1 μg/mL puromycin (+2 μg/mL DOX, +100nM SCH772984) until ready to freeze aliquots.

### Generating barcode sequence:protein variant map

Barcoded variant libraries were digested overnight with SpeI-HF (NEB cat # R3133S) and NheI-HF (NEB cat# R3131S) then submitted to University of Washington PacBio Sequencing Services for further processing. Briefly, PacBio libraries were generated with the SMRTbell Library Prep kit (Pacific Biosciences) and sequenced on a Sequel II SMRT Cell 8M (Pacific Biosciences). Reads were processed using a custom analysis pipeline available at https://github.com/shendurelab/AssemblyByPacBio to identify and link genes with associated barcodes. Enrich2 was used to convert DNA variants into protein coding mutations.

### Abundance assay

The day before transfection MCP-tagged BRaf variant library cells were plated into 4×15cm at ∼45% confluent. The next day, each plate of cells was transfected with 17 μg (pCDNA5) Myc-tdMCP-SNAP + 1.7 μg (pMAX) GFP + 70 μL Fugene HD (Promega cat# E2311) + 935 μL serum-free DMEM. 48 hrs after transfection, cells were trypsinized and combined into two aliquots (2×15cm plates per aliquot). Each aliquot of cells was fully resuspend in 2 mL cold resuspension buffer, which is modRIPA without detergent supplemented with dummy barcode to minimize off-target barcode enrichment (50mM Tris pH=7.6, 150mM NaCl, 10mM NaF, 4mM MgCl_2_, 5% glycerol, Halt protease inhibitor tablet (1 tablet/50mL, Fisher cat# A32965), 2mM PMSF, RiboLock RNase Inhibitor (4uL/mL, Fisher cat#FEREO0382), 100ng/μL total RNA containing dummy barcode). To lyse the resuspended cells, an equal volume of lysis buffer (50mM Tris pH=7.6, 150mM NaCl, 10mM NaF, 4mM MgCl_2_, 5% glycerol, Halt protease inhibitor tablet (1 tablet/50mL, Fisher cat# A32965), 2mM PMSF, RiboLock RNase Inhibitor (4uL/mL, Fisher cat#FEREO0382), 0.2% NP40) was added. Lysates were cleared by centrifugation at 17,000g, 4°C, 10 min then each sample was split into 1.2 mL and 2.8 mL aliquots and combined with 120 μL anti-flag beads (washed, Sigma cat#M8823) or 140 μL anti-myc beads (washed, CST cat#5698S) respectively. Lysates were rotated end-over-end at 4°C for 3 hrs to capture protein. The beads were subsequently washed 3 times with cold wash buffer (50mM Tris pH=7.6, 150mM NaCl, 10mM NaF, 4mM MgCl_2_, 5% glycerol, 0.1% NP40) then resuspended in 75 μL nuclease-free water and 300 μL TRIzol and stored at -20.

### Activity assay

Anti-pErk beads were generated by combining the following components the day before enrichment and rotating end-over-end at 4°C until ready to use: 350 μL protein A beads (CST cat# 73778L) + 350 μL Tris buffered saline + 70 μL anti-pErk antibody (CST cat# 4370L).

To prepare libraries, untagged BRaf variant library cells were plated into two 15 cm dishes (+2 μg/mL DOX, +100nM SCH772984) the day before transfection. On the day of transfection, the media was replaced with fresh media without SCH772984. In activity scans run with overexpressed CRaf, each 15 cm plate was transfected with 17 μg pcDNA5 CRaf-IRES-pErk reporter + 1.7 μg pMAX GFP + 70 μL Fugene HD (Promega cat#E2311) + 950 μL serum-free DMEM. In scans run without overexpressed CRaf, cells were transfected with 2 μg pcDNA5 pErk reporter + 14 μg empty plasmid + 1.6 μg pMAX GFP + 66 μL FugeneHD + 880 μL serum free DMEM.

48 hrs after transfection, each plate of cells was trypsinized, washed once with DPBS, then fully resuspended in 1 mL cold resuspension buffer supplemented with 1% each phosphatase inhibitor cocktails 2 and 3 (Sigma Cat #P5726 and P0044). To lyse the resuspended cells, an equal volume of lysis buffer was added. The lysates were subsequently cleared by centrifugation at 17,000g, 4°C, 10 min then each sample was split into 1.75 mL and 250 μL aliquots. 150 μL of anti-pErk beads or 25 μL anti-flag beads (Sigma cat#M8823) were added to the 1.75 mL and 250 μL aliquots respectively. The lysates were incubated with beads for 4 hrs at 4°C, rotating end-over-end. After enrichment, the beads were washed three times with 1 mL cold wash buffer, resuspended in 75 μL nuclease-free water and 300 μL TRIzol, then stored at -20.

### CRaf interaction assay

TurboID-CRaf was introduced to the tdMCP-BRaf variant library using lentivirus. To generate virus, one 10cm dish of HEK293T cells were transfected with 6 μg transfer plasmid pLJM1 Myc-TurboID-CRaf, EGFP-IRES-BlastR + 3 μg pMDLg/pRRE-Gag & Pol + 1.5 μg pRSV-Rev + 1.5 μg VSV-G envelope + 42 μL Fugene 6 (Promega cat#E2691) + 600 μL serum-free DMEM. 48 hrs after transfection the media was collected and filtered through a 0.4 μm frit. Half of the filtered media was applied to a 15cm dish of the tdMCP-BRaf variant library. Prior to transduction, the variant library was supplemented with RPLP cells stably expressing barcoded Flag-tdMCP-BTK at a frequency of 0.1% of the cell population. BTK barcodes were used as an internal indicator of background labeling and non-specific enrichment. The transduced cell population was maintained for 3 days in complete media (+2 μg/mL DOX, +100nM SCH772984) supplemented with regular FBS. On day 3, the cells were transferred to media containing dialyzed FBS (+DOX, +SCH772984). On day 4 the cells were plated into 2×15cm and 2×10cm dishes, each 50% confluent in media without SCH772984. Drug treatment, biotin labeling and enrichments were performed on day 5.

On day 5, cells were treated with DMSO (0.1% final concentration), 2 μM LXH254, or 5 μM Vemurafenib. After 1 h incubation with drug, in-cell proximity labeling was performed by adding biotin to a final concentration of 200 μM (0.15% DMSO final concentration). The cells were returned to the incubator for 7.5 min at which point the labeling was quenched by trypsinizing the cells, washing once with DPBS, then proceeding immediately to cell lysis. The cells in the 10cm plates were not treated with biotin and were reserved for flag enrichment.

To lyse the cells, each cell pellet was first fully resuspended in 1mL (15 cm plates) or 500 μL (10 cm plates) cold resuspension buffer. To initiate lysis, an equal volume of lysis buffer was added to the resuspended cells. The resulting lysate was cleared by centrifugation at 17,000g, 4°C, for 10 min. To enrich biotin-labeled variants, 2 mL of cleared lysate was combined with 100 μL streptavidin conjugated-beads (Fisher cat# PI20361). 1 mL lysate from unlabeled plates was combined with 100 μL anti-flag beads (Sigma cat#M8823). Samples were rotated end-over-end at 4°C for 1 h then the beads were washed 3 times with 1 mL cold wash buffer. After removing the buffer, the beads were resuspended in 75 μL water and 300 μL TRIzol then stored at -20.

### Mek interaction assay

The day prior to transfection, MCP-tagged BRaf variant library cells were plated into two 15 cm dishes. On the day of transfection cells were washed once with DPBS and plated with fresh media containing DMEM supplemented with 10% dialyzed FBS (Sigma-Aldrich cat# F0392) and 2 μg/mL doxycycline + 100nM SCH772984. This variant library was also supplemented with RPLP cells stably expressing barcoded Flag-tdMCP-BTK at a frequency of 0.1% of the population which was used as an internal metric of background signal. The resulting mixed-population was transfected with 16 μg pcDNA5 TurboID-Mek1 + 1.6 μg pMAX GFP + 66 μL Fugene HD (Promega cat#E2311) + 880 μL DMEM. 24 hrs after transfection, the culture was expanded to two 15 cm plates and two 10 cm plates in DMEM (+dialyzed FBS, +DOX, +100nM SCH772984). 48 hrs after transfection biotin labeling was performed on the 15cm cultures. To label, cells were incubated at 37°C in media supplemented with 250 μM biotin for 5 min. To halt labeling cells were trypsinized, washed once with DPBS and immediately lysed.

To lyse, the collected cells were first fully resuspended in 1 mL (15 cm plates) or 440 μL (10cm plates) resuspension buffer. To lyse the resuspended cells, an equal volume of cold lysis buffer was added. The resulting lysates were cleared by centrifugation at 17,000g, 4°C, 10 min. 100 μL streptavidin beads (washed, Fisher cat# PI20361) was added to the 2 mL of supernatant corresponding to each biotin-labeled replicate. 50 μL anti-flag beads (Sigma cat#M8823) were added to 800 μL of supernatant corresponding to the unlabeled samples. Lysates were rotated end-over-end at 4°C for 45 min then beads were washed 3×1 mL using cold wash buffer. After washing, beads were resuspended in 75 μL nuclease-free water and 300 μL TRIzol then stored at -20.

### SJF0628 degradation assay

The reference, 2xMyc-tdMCP-SNAPtag, was introduced to the tdMCP-BRaf variant library using lentivirus. To generate virus, five 10 cm dishes of HEK293T cells were each transfected with 6 μg transfer plasmid pLJM1 2xMyc-tdMCP-SNAP, EGFP-IRES-BlastR + 3 μg pMDLg/pRRE-Gag & Pol + 1.5 μg pRSV-Rev + 1.5 μg VSV-G envelope + 42 μL Fugene 6 (Promega cat#E2691) + 600 μL serum-free DMEM. 48 hrs after transfection the media from each plate was collected, pooled, and filtered through a 0.4 μm frit. 6% of the filtered media was applied to each of 9×15cm dishes of the tdMCP-BRaf variant library (+2 μg/mL DOX, 100nM SCH772984). 24 hrs after transduction, media was replaced with fresh media without Erk inhibitor (+DOX, NO SCH772984). 48 hrs after transduction, cells were expanded to 12×15cm dishes and treated with SJF0628 (+DOX, NO SCH772984). Final concentrations of SJF0628 were 2.5μM, 500nM, 100nM, 20nM, 4nM, DMSO only and two replicates were performed per drug concentration.

20 hrs after treatment with SJF0628 each plate of cells was lysed. To start, each plate was trypsinized, the media was removed, and cells were fully resuspended in 1 mL cold resuspension buffer. To lyse the resuspended cells, an equal volume of lysis buffer was added. Lysates were cleared by centrifugation at 17,000g, 4°C, 10 min then each sample was split into 540 μL and 1260 μL aliquots and combined with 40 μL anti-flag beads (washed, Sigma cat#M8823) or 120 μL anti-myc beads (washed, CST cat#5698S), respectively. Lysates were rotated end-over-end at 4°C for 3 hrs to capture protein. The beads were subsequently washed 3 times with 1 mL cold wash buffer then resuspended in 75 μL nuclease-free water and 300 μL TRIzol and stored at -20.

### Generating library sequencing amplicons

To prep barcodes for sequencing, RNA was extracted from the enrichment beads using IBI scientific TRI-Isolate RNA Pure Kit (VWR cat# 76006-526) and eluted into 50 μL nuclease-free water. RNA was further concentrated using Zymo RNA Clean & Concentrator (Zymo Research Cat #R1015), eluted into 10 μL nuclease-free water, and stored at -80°C. Following the manufacturer’s instructions, concentrated RNA was reverse transcribed using SuperScript™ IV Reverse Transcriptase (Invitrogen Cat#18090010) and custom primer JJS1 (this manuscript). Briefly, primer/template mixes corresponding to a single library enrichment were generated by combining 7 μL concentrated RNA + 0.625 μL JJS1 (2uM) + 0.625 μL dNTP’s (10mM ea.). The primer was annealed by heating mixes to 75°C for 5 min then set on ice. 0.625 μL DTT (0.1M) + 0.625 μL Ribolock RNase inhibitor (Fisher cat#FEREO0382) + 0.625 μL SSIV enzyme + 2.5 μL SSIV buffer were added to each reaction. cDNA was generated by heating reaction at 45°C for 30 min, 80°C for 10 min, then holding at 4°C.

cDNA was either stored at -20 or immediately used in PCR round #1 to generate double stranded DNA and add amplicon adapters. Briefly, each RT reaction was split into two PCR reactions. Each PCR reaction included 6.25 μL RT reaction + 2.5 μL JJS8 (10uM) + 2.5 μL JJS9-JJS15 (10uM) + 13.75 μL nuclease-free water + 1.5 μL DMSO + 25 μL Q5® Hot Start High-Fidelity 2X Master Mix (NEB cat #M0494). Cycling included 98°C for 30s, five cycles of (98°C for 10s > 64°C for 20s > 72°C for 30s), 72°C for 2 min, and 4°C hold. Replicate reactions were pooled then cleaned with Sera-Mag Select beads (Cytiva #29343052) using a 2.5:1 ratio of beads:reaction and eluted using 20 μL nuclease-free water. Illumina adapters were added in a second round of PCR. This reaction included 5-20 μL of cleaned reaction #1 + 2.5 μL JJS16-JJS23 (10uM) +2.5 μL JJS24 (10uM) + 0.5 μL 10x Sybr gold (Fisher cat#S11494) + 25 μL Q5 MM + 1.0 μL DMSO + nuclease-free water to 51.5uL. PCR cycling included one round at 98°C for 30s, then cycling at (98°C for 10s > 63°C for 20s > 72°C for 30s). Reaction progress was monitored on a BioRad CFX Connect and stopped when RFU reached ∼3,000 (10-15 cycles). Amplicons were pooled based on final RFU values, separated by gel electrophoresis and the band at ∼220bp excised using Freeze ’N Squeeze™ DNA Gel Extraction Spin Columns (Biorad cat#7326166). The DNA was further concentrated using Zymo DNA clean and concentrate columns (Zymo cat#D4014) and quantified using Qubit™ dsDNA HS Assay Kit (Fisher cat#Q32851). Amplicons were sequenced on a NextSeq™ 550 or 2000 using custom primers JJS25-28, 16 cycle paired-end reads and 8 cycle dual indexing.

### Sequencing

Amplicons were sequenced on a NextSeq 500 using a NextSeq 500/550 High Output v2.5 75 cycle kit (Illumina cat #20024906) or on a NextSeq 2000 using a NextSeq 1000/2000 P2 Reagents 100 cycle kit (Illumina cat #20046811). Barcodes were read twice by the paired-end sequencing primers JJS25 and JJS26. Dual indices were sequenced using JJS27 and JJS28. Sequencing reads were converted from BCL to FASTQ format and de-multiplexed using bcl2fastq. Paired sequencing reads were joined by PANDAseq (v2.11) using BFSRKM assembly.

### Calculating variant scores and classifications

Scripts used to filter barcodes and calculate scores are available on the GitHub repository (see URLs section). Frequency values for each barcode in an experiment (F_b_) were generated by calculating the sum of counts recorded for that barcode in all enrichments and dividing by the sum of counts recorded across all barcodes and enrichments.

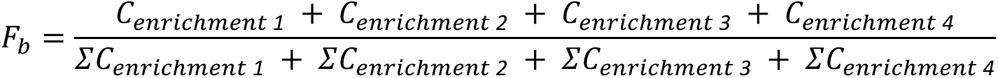

For example, in pErk experiments with two replicates there would be four enrichments including two selective anti-pErk immunoprecipitations and two normalizing anti-flag immunoprecipitations. This frequency calculation was performed for each experiment.

These frequency values were used to filter out low-frequency barcodes which we reasoned would contribute undue levels of counting noise. To determine a frequency cutoff for each experiment, we started with the assumption that accurate synonymous wild type variant scores should form a symmetrical distribution centered at the wild type score. Note, in all experiments wild type was overrepresented with approximately 80 barcodes and therefore the wild score was assumed to be highly accurate. In each experiment we evaluated the impact of a range of minimum F_b_ thresholds on the width and central tendency of the distribution of synonymous wild-type scores by excluding barcodes falling below the specified minimum. The selected threshold F_b_ values, which were unique to each experiment, minimized the skew of the synonymous distribution while retaining most variants.

For pErk enrichments, an additional filter was applied that removed all barcodes containing the motif ‘ATAAA’ or ‘ATTAA’ from our analyses. When compared to other barcodes mapped to the same variant, sequences containing these motifs were disproportionately enriched suggesting they were being directly enriched by the anti-pErk antibody.

To generate variant scores, we first calculated a ratio for each barcode (r_b_) that passed the filters above. For activity scans, the ratio (r_b,act_) was the barcode count observed in the selective pErk enrichment (C_pErk_) normalized to the barcode count observed by enriching all pErk reporter through Flag (C_Flag_).

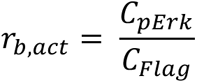

The interaction scan barcode ratio (r_b,int_) was the barcode count observed in the selective streptavidin enrichment (C_Strept_) normalized to the barcode count observed after enriching all variants through Flag (C_Flag_).

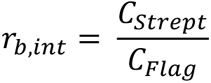

The abundance barcode ratio (r_b,ab_) was the barcode count from the variant enrichment (C_Flag_) divided by the barcode count from the tdMCP standard enrichment (C_Myc_).

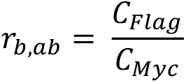

In activity, interaction, and abundance assays the variant score (S_Var_) is the mean r_b_ across all barcodes mapped to the variant (<u>*r*_*b,var*_</u>) normalized to the mean wild type barcode ratio (<u>*r*_*WT*_</u>).

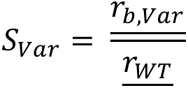

Using these definitions, the wild type score is always set at 1. Scores greater than 1 indicate variants with increased activity, interactions, or abundance. Conversely, scores less than 1 indicate variants with decreased activity, interactions, or abundance.

For each scan the distribution of synonymous wild type variant scores was used to classify each variant. First, we calculated a cutoff value defined as two standard deviations from the mean of the synonymous distribution (2SD_synon_). Next, using the wild type score (1) we defined upper and lower threshold values as

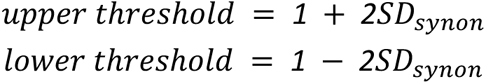

Variants with scores greater than the upper threshold value were classified as ‘increased abundance’, ‘increased activity’, or ‘increased interaction’. Variants with scores lower than the threshold value were classified as ‘decreased abundance’, ‘decreased activity’, or ‘decreased interaction’. Variants with scores that fell between the threshold values were classified as ‘Wild Type-like’.

### Hierarchical clustering

To cluster residues by their biochemical properties, we first transformed into zscores our CRaf interaction scores, Mek interaction scores, activity scores measured with CRaf, CRaf-promoted activity scores, and abundance scores. Residues were then clustered using an average distance calculation and giving equal weight to all variants across all scores.

### Logistic modeling

The growth factor-independent proliferation predictive model was built using the scikit-learn LogisticRegression package and liblinear solver. Model features were z score-transformed LABEL-seq activity scores, CRaf interaction scores, Mek1 interaction scores, abundance scores, and CRaf-promoted activity.

The model built to predict variant sensitivity to SJF028 degradation was created using the scikit-learn LogisticRegression package with the liblinear solver. While training the model, 20% of the data was reserved for model evaluation. Model features were z score-transformed LABEL-seq activity scores, CRaf-promoted activity, CRaf interaction scores, Mek1 interaction scores, abundance scores, Vemurafenib-promoted heterodimerization scores, and LXH254-promoted heterodimerization scores. Predicted values were LABEL-seq SJF0628 degradation classifications.

## Data and code availability

Sequencing data is available in the Sequence Read Archive (SRA) under accession number #########. Scripts written for parsing data and plotting figures are available on Github (http://github.com/MalyLab/########). The code used for PacBio subassembly is also available on Github (https://github.com/shendurelab/AssemblyByPacBio).

## Notes

### Competing Interest Statement

The authors have declared no competing interest.

### Summary of Updates

The figures in this version have been revised.

